# Twin-prime editing enables endogenous protein visualization and mutant allele tracking

**DOI:** 10.64898/2026.09.21.753164

**Authors:** Anghara Menendez, Carlota Ramos, Sebastian Pons

## Abstract

Precise analysis of endogenous protein behavior in mosaic tissues requires strategies that both modify native alleles and identify successfully edited cells at single-cell resolution. Here, we establish a CRISPR-based genome-editing platform that combines endogenous protein visualization with allele-specific mutant tracking in vivo. Using the chick neural tube and β-catenin as a model system, we validate cytosine base editing, prime editing, and twin-prime editing by introducing stabilizing mutations that reproduce their expected cellular and morphological phenotypes. We then develop a twin-prime editing strategy that couples installation of a defined oncogenic mutation to simultaneous insertion of minimal peptide tags, thereby making productive editing directly observable at single-cell resolution without clonal selection. Comparative analysis of ALFA, V5, and split-GFP tags identifies split-GFP as the most reliable strategy for endogenous protein visualization in vivo, and fluorescence-based cell selection strongly enriches for the intended twin-prime editing product. Finally, we extend the approach to endogenous wild-type β-catenin, identifying a functionally neutral insertion site that enables visualization at physiological, non-stabilized levels while preserving normal protein behavior. Together, these results establish twin-prime editing as a versatile platform for directly linking precise endogenous genome modification to protein visualization and allele-specific mutation analysis in intact vertebrate tissues.

## INTRODUCTION

Most vertebrate genetic models, including transgenic mice and embryonic stem cell-derived organoids, do not faithfully reproduce the earliest stages of somatic mutation acquisition. This limitation is particularly important when studying oncogenic mutations, whose biological consequences depend not only on the mutation itself but also on endogenous gene dosage, temporal dynamics and cellular context during tumor initiation. In many experimental systems, mutations are analysed only after prolonged periods of clonal selection, adaptation or tissue expansion, processes that can substantially alter the initial cellular response. Likewise, many cancer models rely on widespread oncogene activation, a scenario fundamentally different from the stochastic mutation of a single cell, frequently affecting only one allele, that characterizes tumor initiation in vivo.

Conventional *in ovo* electroporation has become a powerful approach for studying neural development because it enables rapid, mosaic genetic manipulation and immediate phenotypic analysis in the chick embryo (Muramatsu et al., 1997; Herrera et al., 2014; Herrera et al., 2023). However, most electroporation-based approaches rely on cDNA expression driven by heterologous promoters and intron-less expression cassettes, which do not reproduce endogenous transcriptional regulation, mRNA processing or physiological protein dosage. Precision genome editing combined with *in ovo* electroporation offers an attractive alternative by enabling endogenous mutations to be introduced directly into individual cells while preserving native gene regulation.

The development of CRISPR-based genome editing has progressively expanded the repertoire of programmable genetic modifications that can be introduced into endogenous loci. Base editors enable efficient C•G-to-T•A or A•T-to-G•C conversions without generating double-strand DNA breaks (Komor et al., 2016), whereas prime editors further extend these capabilities by introducing virtually any small substitution, insertion or deletion through reverse transcription of an RNA-encoded template (Anzalone et al., 2022). More recently, engineered pegRNAs (epegRNAs) have improved prime-editing efficiency by stabilizing the guide RNA, and twin-prime editing (twinPE) has enabled substantially larger and more precise sequence insertions through coordinated editing by two complementary prime-editing guide RNAs (Anzalone et al., 2022). Together, these technologies have transformed genome editing from a gene-disruption approach into a versatile platform for precise sequence rewriting.

Despite these advances, a major limitation remains. Whereas therapeutic genome-editing studies typically evaluate editing efficiency at the population level using next-generation sequencing, cell biological studies require the identification of individual edited cells within intact tissues. Current knock-in strategies can introduce fluorescent or epitope tags into endogenous loci, but their low efficiency in the absence of selection makes it difficult to distinguish correctly edited cells from cells carrying unsuccessful editing events or indels (Leonetti et al., 2016; Roberts et al., 2017). Consequently, visualization of endogenous proteins following genome editing remains considerably more challenging than the editing process itself.

An ideal solution would directly couple successful genome editing to protein visualization, so that fluorescence itself reports completion of the intended edit. Such a strategy would permit both the visualization of endogenous proteins and the simultaneous introduction of defined disease-associated mutations during the same editing event. Although prime editing is, in principle, capable of generating these modifications, the insertions required for protein tagging approach the practical limits of editing efficiency and incomplete editing events may still generate detectable signal (Anzalone et al., 2022). Twin-prime editing provides an attractive alternative because the inserted sequence is reconstructed from two independent editing reactions, allowing longer insertions while greatly reducing the probability of false-positive tag generation. These features suggest that twin-prime editing may provide a general framework for selection-free visualization of endogenously edited proteins at single-cell resolution.

Here, we establish CRISPR-based precision genome editing as an experimental platform for endogenous protein visualization and allele-specific analysis in vivo. Using β-catenin as a model system, we first validate cytosine base editing, prime editing, and twin-prime editing in the chick neural tube following *in ovo* electroporation. We then identify split GFP11 as an optimal minimal tag for endogenous protein visualization and develop a twin-prime editing strategy that couples the installation of defined mutations to direct visualization of the edited endogenous protein. Finally, we extend this approach to endogenous, non-stabilized β-catenin, demonstrating single-cell visualization while preserving normal protein behavior. Together, these results establish a versatile strategy for directly linking precise endogenous genome modification to protein visualization and allele-specific functional analysis in vivo.

## RESULTS

### Base editing is highly efficient following in ovo electroporation

We selected cytosine base editing as our initial strategy for in ovo genome editing because of its high efficiency, despite the constraint imposed by its fixed editing window. We used pCMV-CBE6b-SpCas9-D10A-2×UGI (CBE6b), a sixth-generation cytidine base editor with enhanced activity and reduced sequence-context dependence (Zhang et al., 2024), together with pGuia, an sgRNA-expression vector generated in our laboratory for chick neural tube electroporation. pGuia was derived from pSHIN by replacing its H1 promoter with the U6 promoter and sgRNA scaffold from pX330 while retaining the GFP reporter. A GFP-less version, pbGuia, was also generated (Supplementary Fig. 1A).

We targeted the same β-catenin sequence previously used to introduce the stabilizing S33F mutation in mouse embryos (Yeh et al., 2018). HH12 chick embryos were electroporated with CBE6b and pGuia expressing either an sgRNA targeting β-catenin S33 or a scrambled control and analysed 24 or 48 h post-electroporation (hpe; Fig. 1A).

**Figure 1.**
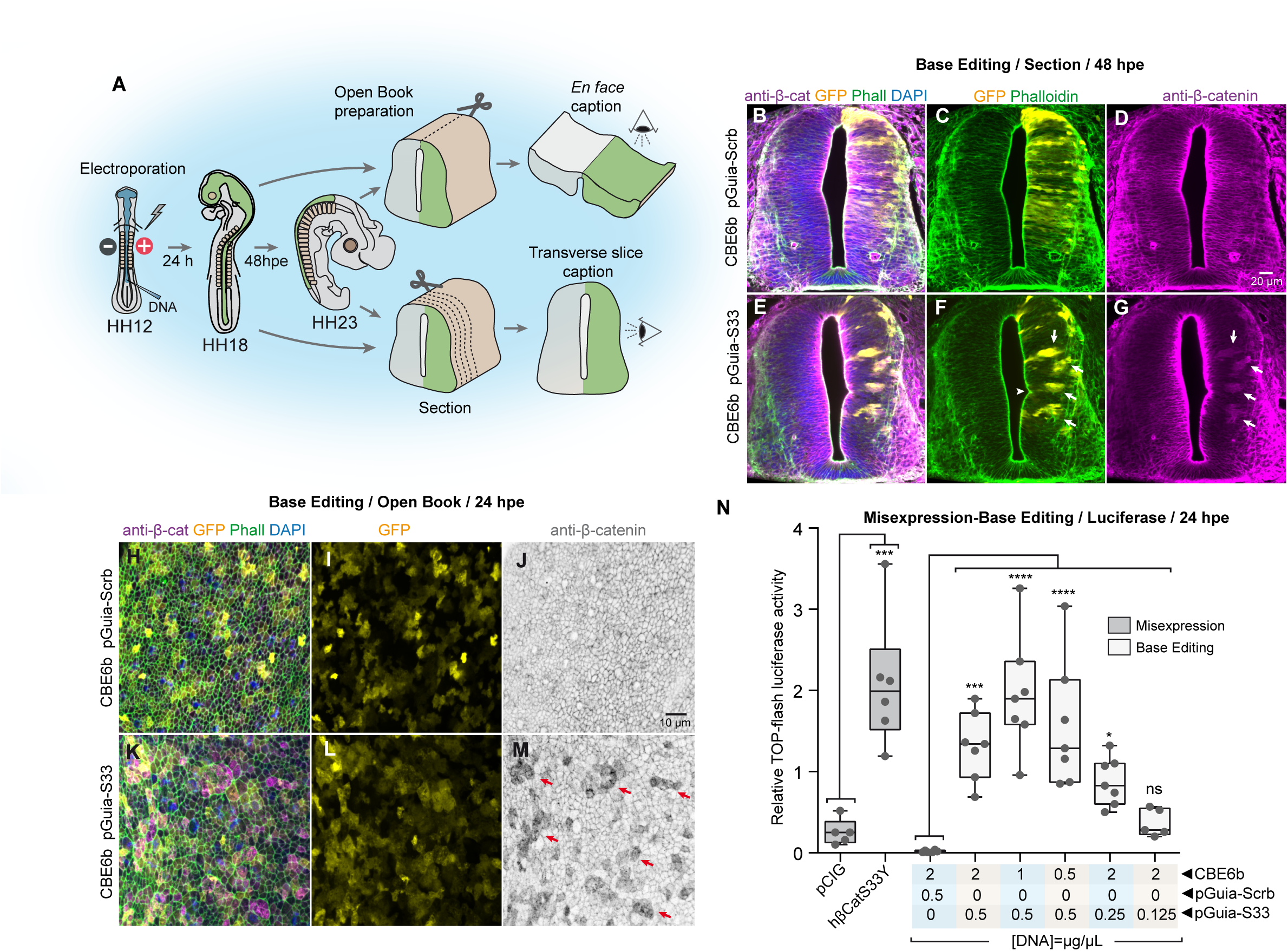
Base editing efficiently induces β-catenin stabilization in the chick neural tube following *in ovo* electroporation. **(A)** Schematic of the experimental workflow. HH12 chick embryos were electroporated *in ovo* and analysed 24 or 48 h post electroporation (hpe). Embryos were processed either as open-book preparations for en face imaging of the apical neuroepithelium or as transverse spinal cord sections. **(B–G)** Transverse sections of chick neural tubes electroporated at HH12 with CBE6b together with pGuia-Scrb (**B–D**) or pGuia-S33 (**E–G**) and analysed 48 hpe. Panels **B** and **E** show merged anti-β-catenin, GFP, phalloidin and DAPI channels. Panels **C** and **F** show the GFP and phalloidin channels, whereas **D** and **G** show the anti-β-catenin channel. Arrowheads indicate apical epithelial intrusions and arrows indicate cytoplasmic β-catenin accumulation. **(H–M)** Open-book preparations of chick neuroepithelium electroporated at HH12 with CBE6b together with pGuia-Scrb (**H–J**) or pGuia-S33 (**K–M**) and analysed 24 hpe. Panels **H** and **K** show merged anti-β-catenin, GFP, phalloidin and DAPI channels. Panels **I** and **L** show the GFP channel alone, whereas panels **J** and **M** show the anti-β-catenin channel. Arrows indicate cells with increased cytoplasmic β-catenin. **(N)** Relative TOP-Flash luciferase activity measured 24 hpe in embryos subjected to base editing using the indicated plasmid DNA concentrations of CBE6b and pGuia-S33 or pGuia-Scrb in the electroporation solution. Misexpression of human β-catenin S33Y from pCIG was included as a positive control. Each point represents an independent embryo. Box-and-whisker plots show the median (center line), the 25th–75th percentiles (box), and the minimum and maximum values (whiskers). Statistical significance is indicated as follows: * *p* < 0.05; *** *p* < 0.001; **** *p* < 0.0001; ns, not significant.

Confocal analysis of transverse spinal cord sections and en face open-book preparations showed β-catenin accumulation specifically in embryos receiving the S33-targeting sgRNA (Fig. 1B–M). Transfected cells formed compact clusters associated with apical epithelial intrusions (Fig. 1F, arrowhead) and showed increased cytoplasmic β-catenin in addition to its junctional pool (Fig. 1G,M, arrows), closely reproducing the phenotype observed following misexpression of stabilized β-catenin S33Y (Herrera et al., 2014) (Supplementary Fig. 1B–E). Because the antibody detects both edited and unedited β-catenin, immunofluorescence did not permit direct quantification of editing efficiency.

We therefore optimized plasmid concentrations using TOP-Flash luciferase activity as a functional readout. Maximal activation was obtained with 0.5 μg/μL pGuia and 1 μg/μL CBE6b, reaching levels comparable to those induced by pCIG-driven β-catenin S33Y expression (Fig. 1N).

To confirm editing at the endogenous locus, H2B-RFP-positive cells were isolated by FACS and subjected to Oxford Nanopore amplicon sequencing of ggCTNB1 exon 4 (Supplementary Fig. 2A). Overall, 71.34% of reads corresponded to WT and 28.66% contained sequence variations. The two most abundant edited alleles represented 7.56% and 2.68% of total reads. Both contained the C•G-to-T•A substitution generating S33F, with the former carrying an additional synonymous substitution at −11. Other variants were individually less frequent and heterogeneous, and a contribution from the intrinsic error rate of Oxford Nanopore sequencing cannot be excluded. Thus, sequencing directly confirmed installation of the intended endogenous S33F mutation, consistent with the strong Wnt/β-catenin activation observed in vivo.

### Prime editing enables precise β-catenin S33 mutagenesis in the chick neural tube

The fixed editing window of base editors restricts the mutations that can be introduced. We therefore evaluated prime editing as a more versatile strategy. We implemented PE3 using PEmax and epegRNAs expressed from pU6-tevopreQ1-GG, together with a second-nick sgRNA positioned 55 bp downstream of the primary nick (Fig. 2A–C). The S33Y epegRNA contained a 20-nt spacer, 13-nt PBS, 16-nt reverse-transcription template and tevopreQ1 stabilization motif.

**Figure 2.**
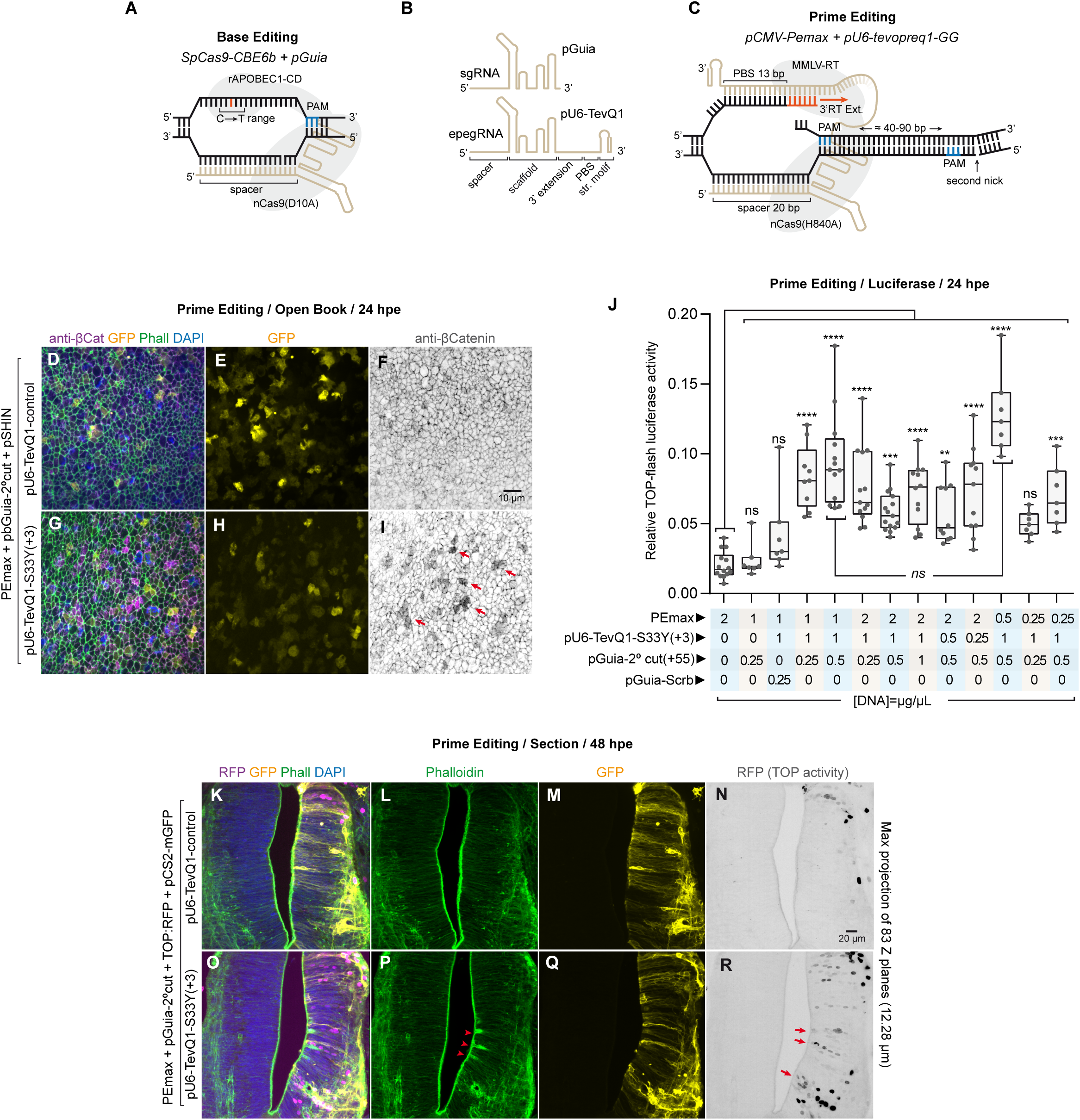
Prime editing enables precise installation of the β-catenin S33Y mutation in the chick neural tube. **(A–C)** Schematic comparison of cytosine base editing and prime editing. Cytosine base editing is constrained by a fixed editing window relative to the PAM, whereas prime editing enables precise sequence modification through reverse transcription. Panel **B** shows a schematic comparison of a canonical sgRNA and an engineered prime editing guide RNA (epegRNA). The epegRNA contains a primer binding site (PBS), reverse transcription (RT) template and a structured RNA motif that stabilizes the 3′ extension. Panel **C** shows the PE3 strategy used in this study. PEmax consists of a Cas9(H840A) nickase fused to an MMLV reverse transcriptase and guided by an epegRNA expressed from pU6-tevopreQ1-GG. A second nick was introduced on the non-edited strand using pGuia-2°cut positioned 55 bp downstream of the primary nick. **(D–I)** Open-book preparations of chick neural tubes electroporated at HH12 with PEmax, pGuia-2°cut, pSHIN and either a control epegRNA (**D–F**) or the β-catenin S33Y epegRNA (**G–I**) and analysed 24 hpe. Panels **D** and **G** show merged anti-β-catenin, GFP, phalloidin and DAPI channels. Panels **E** and **H** show the GFP channel alone, whereas **F** and **I** show the anti-β-catenin channel. Arrows indicate cells with increased cytoplasmic β-catenin. **(J)** Relative TOP-Flash luciferase activity measured 24 hpe in embryos subjected to prime editing using the indicated plasmid DNA concentrations of PEmax, pU6-tevopreQ1-S33Y(+3), pGuia-2°cut and pGuia-Scrb in the electroporation solution. Each point represents an independent embryo. Box-and-whisker plots show the median (center line), the 25th–75th percentiles (box), and the minimum and maximum values (whiskers). Statistical significance is indicated as follows: ** *p* < 0.01; *** *p* < 0.001; **** *p* < 0.0001; ns, not significant. **(K–R)** Maximum-intensity projections generated from 83 confocal Z-planes spanning 12.28 μm of transverse sections of chick neural tubes electroporated at HH12 with PEmax, pGuia-2°cut, TOP-RFP and pCS2-mGFP together with either a control epegRNA (**K–N**) or the β-catenin S33Y epegRNA (**O–R**) and analysed 48 hpe. Panels **K** and **O** show merged RFP, GFP, phalloidin and DAPI channels. Panels **L** and **P** show the phalloidin channel, **M** and **Q** the GFP channel, and **N** and **R** the TOP-RFP channel. Arrowheads indicate apical epithelial intrusions and arrows indicate ectopic TOP-RFP-positive cells.

As an initial functional validation, HH12 embryos were electroporated with PEmax, the S33Y epegRNA and second-nick sgRNA and analysed after 24 h. Open-book preparations showed increased cytoplasmic β-catenin in a subset of cells receiving the S33Y epegRNA (Fig. 2I, arrows), whereas β-catenin remained predominantly junctional in controls (Fig. 2F).

We next optimized the relative concentrations of PEmax, epegRNA and second-nick sgRNA using TOP-Flash. The highest reporter activities were obtained at concentrations of 1–1–0.25, 1–1–0.5 or 0.5–1–0.5 μg/μL, respectively (Fig. 2J). Using the 1–1–0.25 μg/μL combination, embryos analysed after 48 h showed ectopic TOP-RFP-positive cells outside its endogenous dorsal domain (Fig. 2R, arrows), frequently associated with epithelial distortions and apical intrusions characteristic of stabilized β-catenin signaling (Fig. 2P, arrowheads). No ectopic TOP-RFP activation was detected in controls (Fig. 2N).

Amplicon sequencing of H2B-RFP-positive cells provided direct molecular confirmation of the edit (Supplementary Fig. 2B). Of the reads, 77.11% corresponded to WT and 22.89% contained sequence variations. The most abundant non-WT allele (7.38%; 347 reads) carried the intended C•G-to-A•T substitution generating S33Y. Other variants were individually below 1% and heterogeneous, and their interpretation is subject to the intrinsic error rate of Oxford Nanopore sequencing. Thus, prime editing precisely installed S33Y at the endogenous ggCTNB1 locus. However, neither immunofluorescence, reporter activation nor bulk sequencing directly identifies the individual cells carrying the intended edit.

### Selection of an optimal tag for genome editing

To identify edited cells directly, we sought to couple mutation installation to insertion of a minimal detection tag. We compared split GFP11 (Kamiyama et al., 2016), ALFA (Götzke et al., 2019) and V5 (Zeghal et al., 2023). ALFA and V5 were detected using fluorescent chromobodies, whereas GFP11 was visualized by complementation with GFP1–10.

Tag performance was first compared independently of editing efficiency using pTest, a CAG-driven β-catenin S33Y-miRFP670 reporter containing a Golden Gate cassette between residues Y33 and G34 (Fig. 3A). This allowed tag-dependent fluorescence to be compared directly with intrinsic β-catenin-miRFP670 fluorescence. We tested one or three copies of ALFA and V5 and one, two or three copies of GFP11 (Fig. 3B–D).

**Figure 3.**
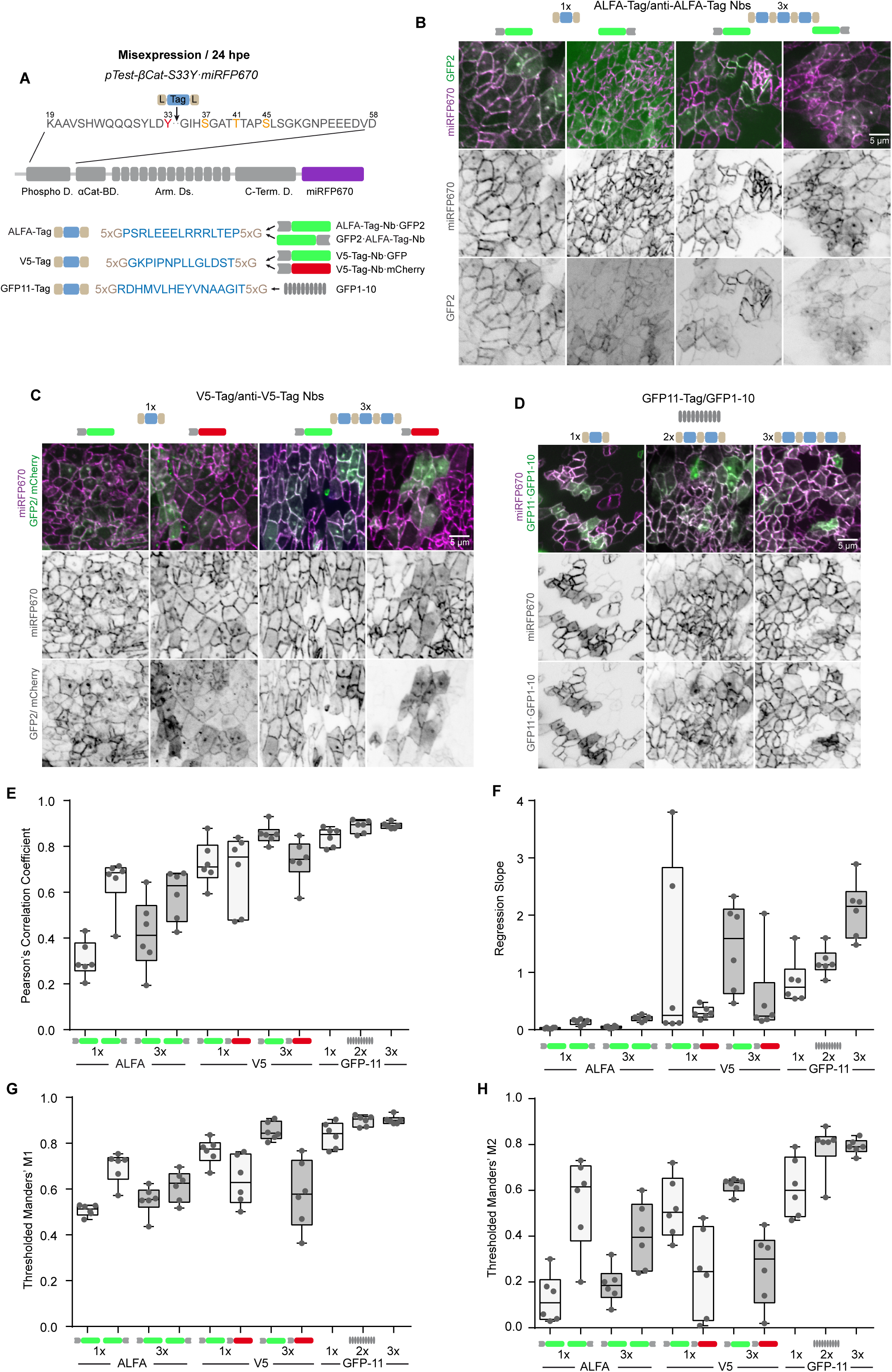
Selection of an optimal tag for endogenous β-catenin genome editing. **(A)** Schematic of the pTest-βCat-S33Y-miRFP670 reporter used to compare candidate tags independently of genome editing efficiency. A Golden Gate cloning cassette (BsmBI–BsmBI) was inserted between residues Y33 and G34 of β-catenin S33Y, allowing modular insertion of ALFA, V5 or GFP11 tags flanked by 5×G linkers. ALFA and V5 were detected using fluorescent chromobodies, whereas GFP11 was detected by complementation with GFP1–10. **(B)** Representative en face confocal images of chick neural tubes electroporated at HH12 with pTest constructs carrying one or three ALFA-tag repeats together with fluorescent ALFA chromobodies and analysed 24 hpe. Merged images show intrinsic miRFP670 fluorescence and GFP2 fluorescence derived from the chromobodies. Individual miRFP670 and GFP2 channels are shown below. **(C)** Representative en face confocal images of chick neural tubes electroporated at HH12 with pTest constructs carrying one or three V5-tag repeats together with GFP- or mCherry-tagged anti-V5 chromobodies and analysed 24 hpe. Merged images show intrinsic miRFP670 fluorescence together with chromobody fluorescence. Individual miRFP670 and chromobody channels are shown below. **(D)** Representative en face confocal images of chick neural tubes electroporated at HH12 with pTest constructs carrying one, two or three GFP11 repeats together with GFP1–10 and analysed 24 hpe. Merged images show intrinsic miRFP670 fluorescence together with GFP complementation, whereas individual miRFP670 and GFP channels are shown below. **(E–H)** Quantitative colocalization analysis comparing tag-dependent fluorescence with intrinsic miRFP670 fluorescence. **(E)** Pearson’s correlation coefficient. **(F)** Regression slope. **(G)** Thresholded Manders’ M1 coefficient. **(H)** Thresholded Manders’ M2 coefficient. Six independent embryos were analysed per condition, and each point represents the value obtained from one embryo. Box-and-whisker plots show the median (center line), the 25th–75th percentiles (box), and the minimum and maximum values (whiskers).

Split GFP most faithfully reproduced β-catenin-miRFP670 localization and showed virtually no detectable background, with fluorescence increasing progressively with GFP11 copy number. ALFA generated the weakest signal, whereas V5 performed better but, like ALFA, showed background and intracellular aggregates in highly expressing cells.

Quantitative colocalization analysis confirmed these observations (Fig. 3E–H). Across Pearson’s correlation, intensity regression slope and thresholded Manders’ coefficients M1 and M2, GFP11 showed the best spatial agreement, proportionality, detection coverage and specificity. GFP11 was therefore selected as the primary tag for endogenous β-catenin editing, with V5 retained as an orthogonal detection tag.

### Twin-prime editing enables simultaneous mutation and visualization of β-catenin

We next asked whether GFP11 could be introduced together with S33Y in a single editing event. A GFP11 tag flanked by two 5×G linkers requires an insertion of approximately 100 bp, approaching the practical range of twin-prime rather than conventional prime editing. We therefore designed one forward epegRNA and two alternative reverse epegRNAs, generating 98-bp (Edit-1) and 104-bp (Edit-2) edits (Fig. 4A,B). The forward epegRNA encoded S33Y and the 5′ portion of GFP11, whereas the reverse epegRNAs encoded the remaining tag sequence; the reverse-transcription products shared a 21-bp overlap. Because S33Y and GFP11 are reconstructed within the same editing event, GFP complementation provides a direct functional readout of productive editing.

**Figure 4.**
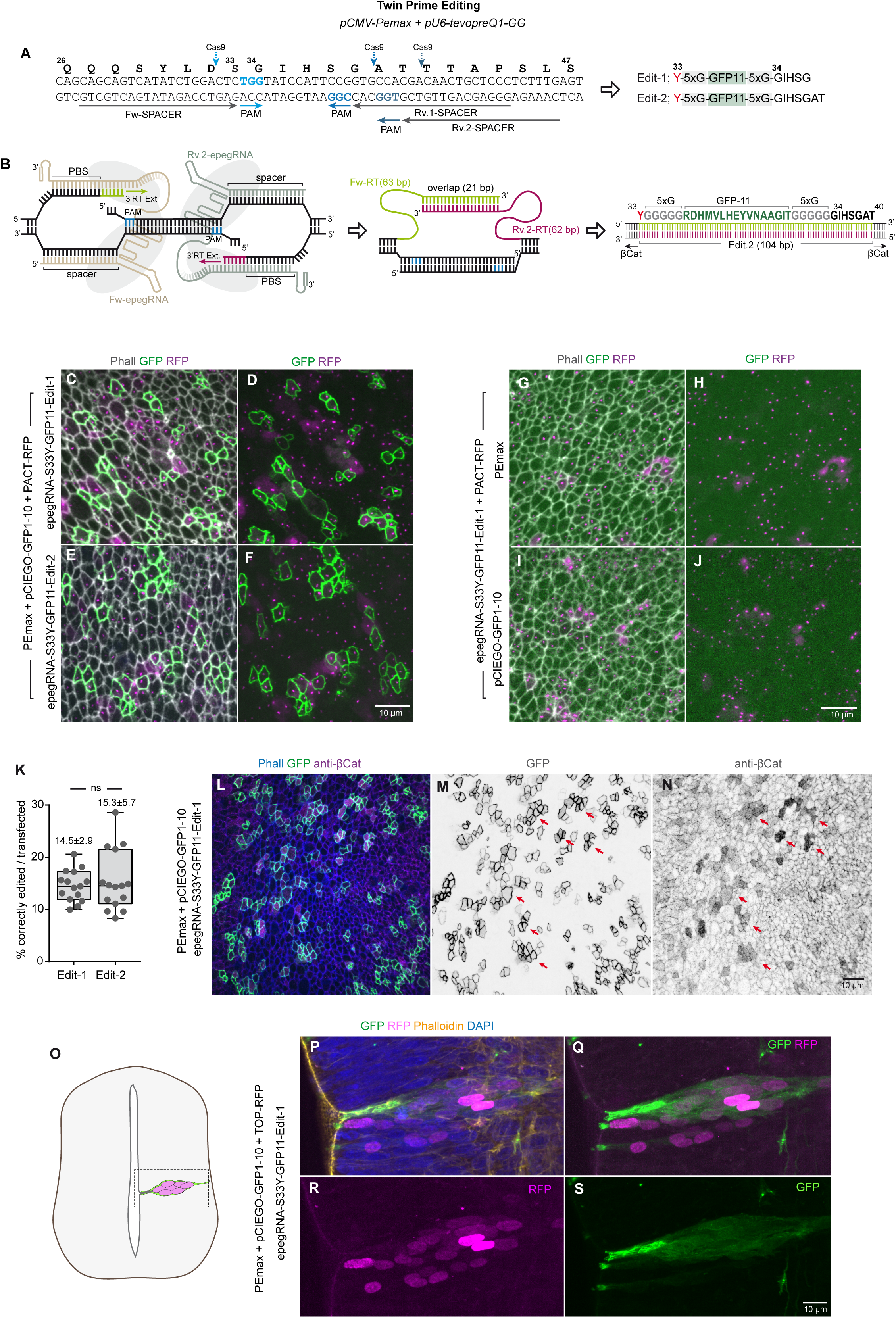
Twin-prime editing enables simultaneous installation of the S33Y mutation and visualization of endogenous β-catenin. **(A)** Schematic of the twin-prime editing strategy used to introduce the S33Y mutation together with a GFP11 tag flanked by 5×G linkers between residues Y33 and G34 of β-catenin. One forward epegRNA and two alternative reverse epegRNAs (Rv.1 and Rv.2) were designed to generate Edit-1 and Edit-2, differing in PAM spacing and final edit size. **(B)** Schematic of the twin-prime editing configuration. Forward and reverse epegRNAs generated overlapping reverse-transcription products encoding complementary portions of the GFP11 insertion. Spacer and primer-binding site lengths were maintained at 20 bp and 13 bp, respectively. Edit-1 and Edit-2 generated final edit sizes of 98 bp and 104 bp, respectively, with a 21-bp overlap between reverse-transcription products. **(C–F)** Open-book preparations of chick neural tubes electroporated at HH12 with PEmax, pCIEGO-GFP1–10, PACT-RFP and either the Edit-1 (**C,D**) or Edit-2 (**E,F**) epegRNA pair and analysed 24 hpe. Panels **C** and **E** show merged phalloidin, GFP and RFP channels. Panels **D** and **F** show the GFP and RFP channels. **(G–J)** Background controls for split-GFP complementation using the Edit-1 configuration. Embryos were electroporated with the Edit-1 epegRNAs together with GFP1–10 but without PEmax (**G,H**), or together with PEmax but without GFP1–10 (**I,J**). Panels **G** and **I** show merged phalloidin, GFP and RFP channels, whereas **H** and **J** show the GFP and RFP channels. **(K)** Quantification of correctly edited cells identified by GFP11 complementation following Edit-1 or Edit-2 twin-prime editing. GFP11-positive cells are expressed as a percentage of PACT-RFP-positive transfected cells. Fifteen fields from six independent embryos were analysed per condition, with each point representing the percentage calculated for one 104 × 104 μm field. Box-and-whisker plots show the median (center line), the 25th–75th percentiles (box), and the minimum and maximum values (whiskers). Values above the plots indicate mean ± SD. Statistical significance is indicated as follows: ns, not significant. **(L–N)** Open-book preparations of chick neural tubes electroporated at HH12 with PEmax, pCIEGO-GFP1–10 and the Edit-1 epegRNA pair and analysed 24 hpe, followed by anti-β-catenin immunostaining. Panel **L** shows merged phalloidin, GFP and anti-β-catenin channels. Panels **M** and **N** show the GFP and anti-β-catenin channels separately. Arrows indicate GFP-positive cells with increased cytoplasmic β-catenin. **(O)** Schematic indicating the spinal cord region analysed in transverse sections. **(P–S)** Transverse sections of chick embryos electroporated at HH12 with PEmax, pCIEGO-GFP1–10, TOP-RFP and the Edit-1 epegRNA pair and analysed 48 hpe. Panel **P** shows merged GFP, RFP, phalloidin and DAPI channels. Panels **Q** and **R** show the GFP/RFP merge and the RFP channel, respectively, whereas **S** shows the GFP channel.

HH12 embryos were electroporated with PEmax, GFP1–10, PACT-RFP and either epegRNA pair and analysed after 24 h. GFP fluorescence localized predominantly to apical adherens junctions, reproducing the expected distribution of β-catenin (Fig. 4C– F). GFP-positive cells represented 14.5 ± 2.9% and 15.3 ± 5.7% of PACT-RFP-positive cells for Edit-1 and Edit-2, respectively (Fig. 4K), indicating no measurable effect of the differences in edit size or PAM spacing. No junctional GFP signal was detected when either PEmax or GFP1–10 was omitted (Fig. 4G–J).

GFP-positive cells also showed elevated cytoplasmic β-catenin by immunostaining (Fig. 4L–N). At 48 h, GFP-positive clusters displayed junctional, cytoplasmic and nuclear β-catenin, coincident TOP-RFP activation and epithelial distortions characteristic of stabilized β-catenin signaling (Fig. 4P–S). An equivalent S33Y-V5 edit was also detectable using fluorescent chromobodies or anti-V5 immunostaining (Supplementary Fig. 1F–L), although its fluorescence was less sharply defined than split GFP.

Amplicon sequencing independently confirmed the twin-prime editing product (Supplementary Fig. 2C,D). The complete S33Y–5×G–GFP11–5×G allele increased from 0.05% of reads in H2B-RFP-selected cells to approximately 6% following GFP-based selection. Additional reads contained GFP11-derived sequences with sequence variations and were analysed separately. Thus, GFP complementation strongly enriched the population for the intended twin-prime editing product.

### epegRNA competition enables estimation of biallelic twin-prime editing frequencies

We next used competitive twin-prime editing to estimate how frequently both β-catenin alleles were edited in the same cell. Two epegRNA pairs targeting identical genomic positions introduced either S33Y-GFP11 or S33Y-V5 (Fig. 5A,B). Because productive tags require homologous forward and reverse epegRNAs, GFP11/V5 double-positive cells report independent installation of different edits on the two β-catenin alleles (Fig. 5C), whereas single-positive cells may be monoallelic or carry the same tag on both alleles.

**Figure 5.**
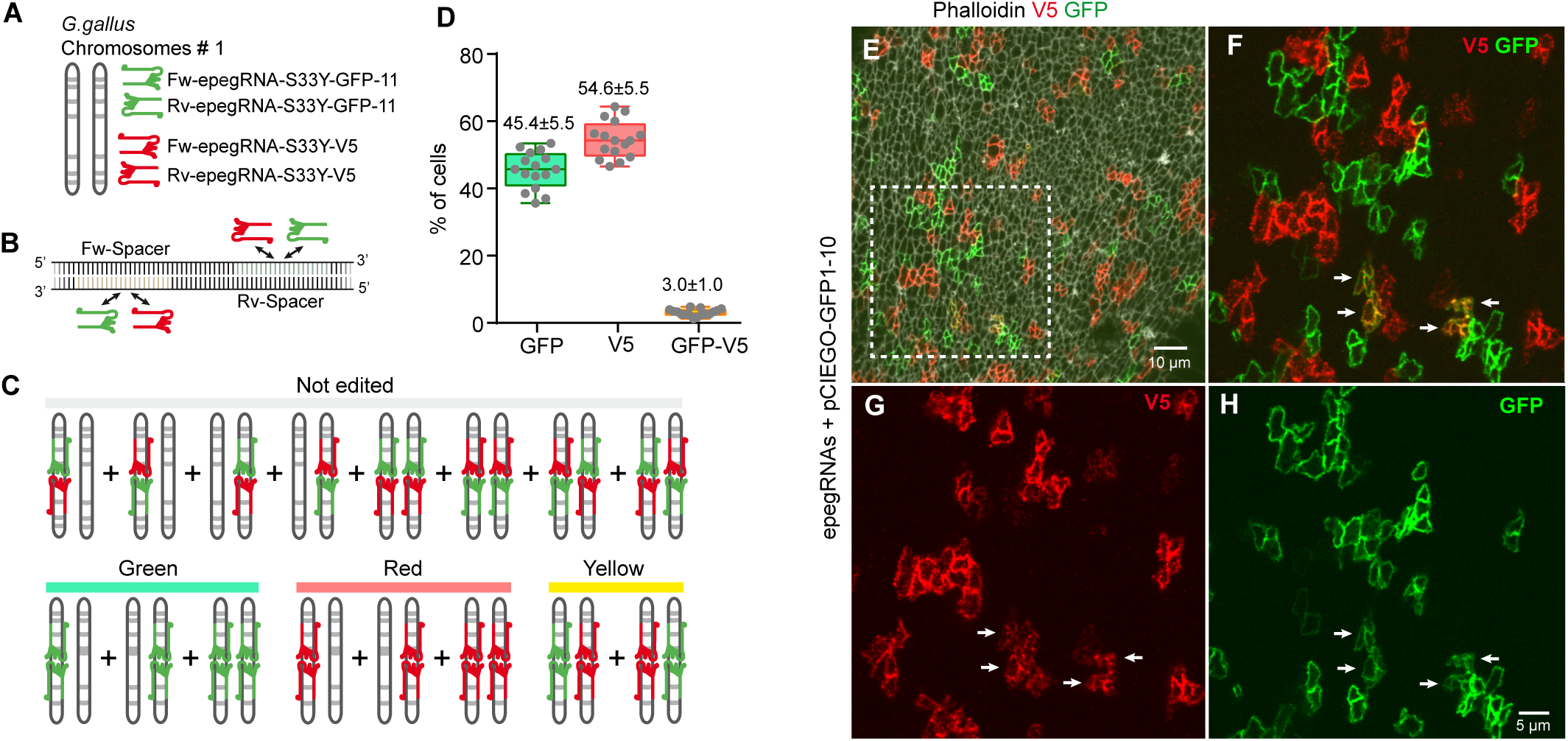
epegRNA competition enables estimation of biallelic twin-prime editing frequencies. **(A)** Schematic of the competitive twin-prime editing strategy. Two independent epegRNA pairs targeting the same forward and reverse spacer sequences were designed to introduce either S33Y-GFP11 or S33Y-V5 at the same position within the endogenous β-catenin locus. **(B)** Diagram illustrating competition between GFP11- and V5-encoding epegRNAs for the same forward and reverse target sites. Productive editing occurs only when homologous forward and reverse epegRNAs act together, generating either GFP11 or V5 insertions. **(C)** Schematic of the possible editing outcomes generated by competitive epegRNA pairing. GFP11-positive cells are shown in green, V5-positive cells in red, double-positive cells in yellow, and non-detectable editing outcomes are indicated separately. **(D)** Quantification of GFP11-positive, V5-positive and GFP11/V5 double-positive cells in chick embryos electroporated at HH12 with all four epegRNAs together with PEmax and pCIEGO-GFP1–10 and analysed 24 hpe. Six independent embryos were analysed, comprising a total of 16 fields (104.6 × 104.8 μm), with each point representing the percentage calculated for one field. GFP11- and V5-positive cells are expressed relative to the combined number of GFP11- and V5-positive cells, whereas double-positive cells were quantified separately using the same denominator. Box-and-whisker plots show the median (center line), the 25th–75th percentiles (box), and the minimum and maximum values (whiskers). Values above the plots indicate mean ± SD. **(E–H)** Representative open-book preparations of chick neural tubes electroporated at HH12 with all four epegRNAs and analysed 24 hpe. Panel **E** shows merged phalloidin, V5 and GFP channels. Panel **F** shows a higher magnification of the boxed region in **E**. Panels **G** and **H** show the V5 and GFP channels separately. Arrows indicate GFP11/V5 double-positive cells.

Among detectable edited cells, 45.4 ± 5.5% were GFP11-positive, 54.6 ± 5.5% V5-positive and 3.0 ± 1.0% double-positive (Fig. 5D–H). The assay cannot distinguish monoallelic editing from biallelic installation of the same tag, and some heterologous epegRNA combinations remain undetectable. Nevertheless, the low frequency of GFP11/V5 double-positive cells indicates that independent biallelic editing is infrequent under these conditions, consistent with a predominance of monoallelic editing.

### Twin-prime editing enables visualization of endogenous β-catenin at single-cell resolution

Because the preceding experiments used stabilized β-catenin, we next asked whether split GFP could visualize non-stabilized endogenous protein. Two twin-prime strategies inserted GFP11 within the N-terminal phosphorylation domain (Fig. 6A,B). WT reproduced the S33Y insertion site between S33 and G34, within the β-TrCP phosphodegron, whereas WT-SP inserted GFP11 between L31 and D32, immediately upstream of the degron and preserving its recognition sequence.

**Figure 6.**
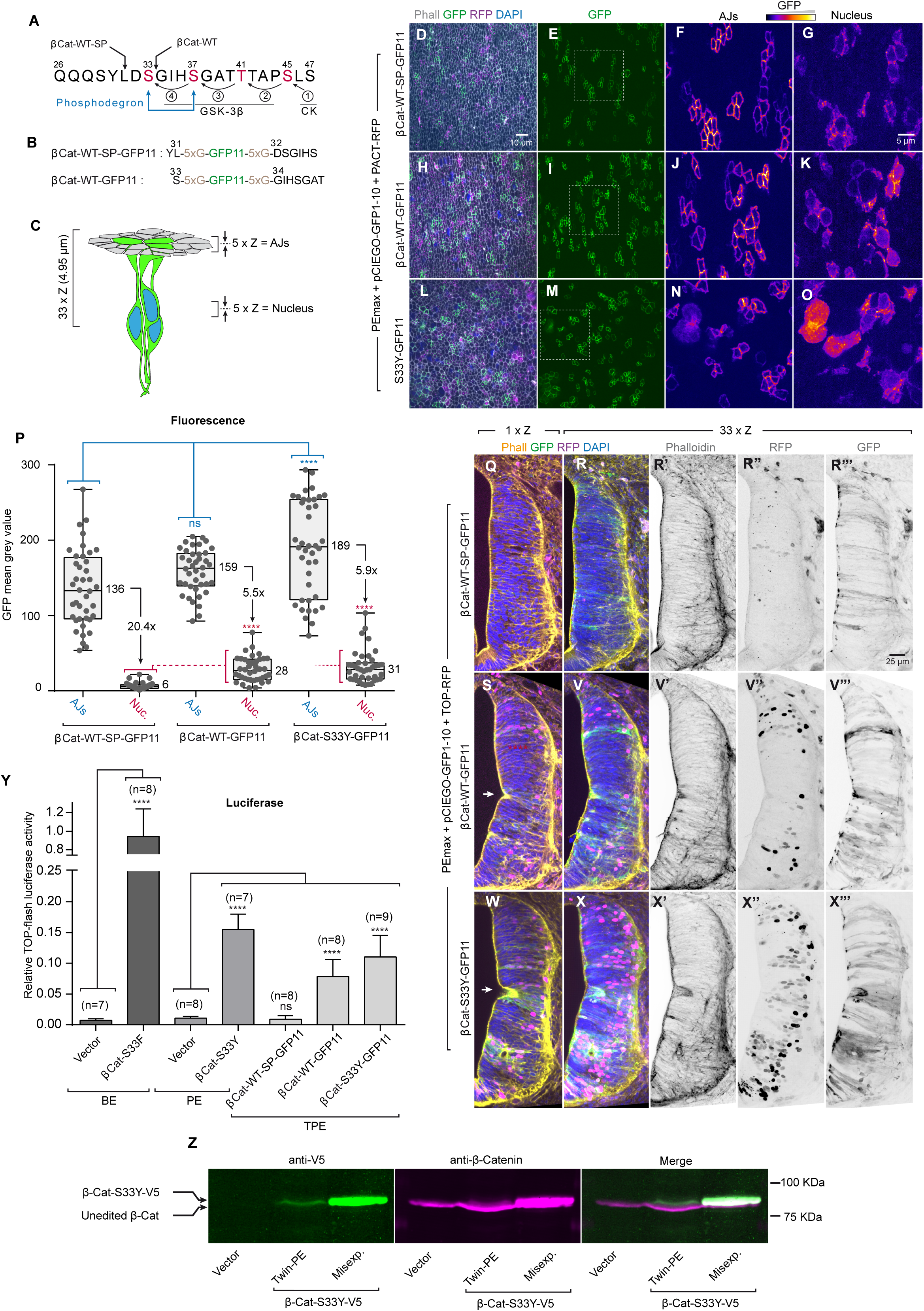
Twin-prime editing enables visualization of endogenous β-catenin at single-cell resolution. **(A)** Sequence of the β-catenin N-terminal phosphorylation domain showing the residues phosphorylated by CK1 and GSK-3β, the β-TrCP phosphodegron motif, and the two GFP11 insertion sites. WT-SP (safe place) and WT denote GFP11 insertion between residues L31 and D32, and S33 and G34, respectively. **(B)** Amino acid sequences generated by the WT-SP-GFP11 and WT-GFP11 edits. GFP11 was flanked by 5×G linkers in both configurations. **(C)** Schematic of the imaging strategy used for quantitative analysis. Five-plane maximum-intensity projections centred on either the apical adherens junctions (AJs) or the nuclear layer were generated from 33-plane confocal stacks spanning 4.95 μm. **(D–O)** Open-book preparations of chick neural tubes electroporated at HH12 with PEmax, pCIEGO-GFP1–10 and PACT-RFP together with WT-SP-GFP11 (**D–G**), WT-GFP11 (**H–K**) or S33Y-GFP11 (**L–O**) and analysed 24 hpe. Panels **D**, **H** and **L** show merged phalloidin, GFP, RFP and DAPI channels. Panels **E**, **I** and **M** show the GFP channel. Panels **F**, **J** and **N** show five-plane maximum-intensity projections centred on the apical adherens junctions, whereas **G**, **K** and **O** show equivalent projections centred on the nuclear layer. **(P)** Quantification of GFP mean gray values at adherens junctions (AJs) and nuclei (Nuc.) in WT-SP-GFP11, WT-GFP11 and S33Y-GFP11 edited cells. Four independent embryos were analysed per condition, with ten measurements obtained from each embryo, yielding 40 measurements per condition. Each point represents an individual AJ or nuclear measurement. Box-and-whisker plots show the median (center line), the 25th–75th percentiles (box), and the minimum and maximum values (whiskers). Values beside the plots indicate mean GFP fluorescence, and the AJ-to-nuclear mean fluorescence ratio is shown for each condition. For statistical analysis, measurements were averaged per embryo, and embryo means (***n*** = 4 per condition) were compared separately for AJs and nuclei using one-way ANOVA followed by Dunnett’s multiple-comparisons test. Statistical significance is indicated as follows: **** *p* < 0.0001; ns, not significant. **(Q–X’’’)** Transverse sections of chick neural tubes electroporated at HH12 with PEmax, pCIEGO-GFP1–10 and TOP-RFP together with WT-SP-GFP11 (**Q–R’’’**), WT-GFP11 (**S–V’’’**) or S33Y-GFP11 (**W–X’’’**) and analysed 48 hpe. Panels **Q**, **S** and **W** show representative single confocal planes including phalloidin, GFP, RFP and DAPI. Panels **R**, **V** and **X** show 33-plane maximum-intensity projections of the same regions, whereas **R’–R’’’**, **V’–V’’’** and **X’–X’’’** show the corresponding phalloidin, RFP and GFP channels. Arrows indicate epithelial intrusions. **(Y)** Relative TOP-Flash luciferase activity measured 24 h after electroporation following β-catenin S33F base editing (BE), S33Y prime editing (PE), or twin-prime editing (TPE) generating S33Y-GFP11, WT-SP-GFP11 or WT-GFP11, together with the corresponding vector controls. Bars represent mean ± SD, and the number of independent embryos analysed (*n*) is indicated above each bar. Statistical significance is indicated as follows: **** *p* < 0.0001; ns, not significant. **(Z)** Western blot analysis of embryos subjected to twin-prime editing or conventional misexpression of βCat-S33Y-V5. Anti-V5 and anti-β-catenin immunoblotting detected V5-tagged β-catenin at the expected molecular weight following both twin-prime editing and conventional misexpression.

In open-book preparations, junctional GFP fluorescence was broadly similar among WT-SP, WT and S33Y, whereas nuclear fluorescence was markedly higher in WT and S33Y than in WT-SP (Fig. 6C–O). Mean junctional fluorescence was 136 ± 8, 159 ± 4.6 and 189 ± 10.1, respectively, with no significant difference between WT-SP and WT. Nuclear fluorescence increased from 6.6 ± 0.6 in WT-SP to 28.6 ± 2.5 in WT and 31.8 ± 3.3 in S33Y, reducing the junctional-to-nuclear ratio from 20.4:1 to 5.5:1 and 5.9:1, respectively (Fig. 6P). Thus, insertion within the phosphodegron partially stabilizes β-catenin, whereas WT-SP preserves its predominantly junctional distribution.

Consistent with this conclusion, WT-SP-edited cells showed no detectable ectopic TOP-RFP activation or tissue deformation (Fig. 6Q–R′′′). By contrast, WT and S33Y showed increased cytoplasmic and nuclear β-catenin, ectopic TOP-RFP activation and epithelial intrusions (Fig. 6S–X′′′).

TOP-Flash assays comparing all editing strategies showed the strongest Wnt activation with S33F base editing, whereas S33Y prime and twin-prime editing produced similar lower activation (Fig. 6Y). WT-SP did not activate the pathway above control levels, whereas WT retained partial stabilizing activity. Twin-prime editing additionally permitted direct identification of edited cells through GFP11 complementation.

Finally, S33Y-V5 β-catenin generated by twin-prime editing was detected at the expected molecular weight by both anti-V5 and anti-β-catenin antibodies (Fig. 6Z). Its abundance was substantially lower than following CAG-driven S33Y-V5 misexpression, consistent with endogenous expression and the lower signaling output observed in the TOP-Flash assays. Together, these experiments establish that twin-prime editing combined with split GFP is sufficiently sensitive to visualize non-stabilized endogenous β-catenin at single-cell resolution and identify WT-SP as a functionally neutral insertion site for comparative β-catenin editing.

## DISCUSSION

Precision genome editing has transformed the ability to introduce defined mutations into endogenous loci, but its application to cell biology presents an additional challenge: identifying which individual cells carry the intended edit and determining how the resulting protein behaves within its native tissue context. This is particularly relevant in mosaic in vivo models, where edited cells coexist with unedited cells and cells carrying incomplete or unintended editing outcomes. Here, we address this limitation by coupling precision genome editing to visualization of the edited endogenous protein. Using the chick neural tube and β-catenin as a model, we establish a workflow linking installation of defined mutations to identification of edited cells, protein localization, pathway activation and tissue phenotype.

The chick embryo is particularly suited to this approach. In ovo electroporation permits rapid and spatially restricted genetic manipulation without stable transgenesis or clonal expansion, but conventional experiments generally rely on cDNAs expressed from heterologous promoters. Although highly informative, such approaches do not reproduce endogenous transcriptional regulation or protein dosage, an important limitation for proteins such as β-catenin, whose activity depends on tightly controlled abundance and subcellular distribution. Combining in ovo electroporation with base, prime and twin-prime editing therefore extends the chick neural tube from a model of mosaic gene misexpression to one in which endogenous alleles can be rewritten and analysed directly.

β-Catenin provides a stringent test of this framework because stabilizing mutations within its N-terminal phosphodegron generate several independent readouts, including protein accumulation and redistribution, Wnt pathway activation and epithelial deformation. Introduction of S33 substitutions by the three editing approaches reproduced these expected effects, supporting the biological relevance of endogenous editing in this system. The comparison also illustrates complementary strengths of the different platforms. Base editing produced the strongest signaling output and is highly effective when the desired substitution is compatible with its editing window. Prime editing expands the range of possible sequence changes, but neither phenotypic readouts nor population-level sequencing readily identify individual cells carrying the intended allele. In this respect, the principal advantage of the twin-prime editing strategy developed here is not simply its capacity to introduce a larger sequence, but its ability to make productive editing directly observable.

This property relies on coupling the mutation to a minimal detection tag within the same editing event. Comparison of ALFA, V5 and GFP11 showed that tag size alone does not determine suitability for endogenous protein imaging; the detection mechanism is equally important. Chromobody-based detection of ALFA and V5 produced background associated with freely diffusing or aggregated fluorescent nanobodies. Split GFP avoided this limitation because fluorescence required complementation between GFP11 inserted into β-catenin and GFP1–10 supplied in trans. GFP11 consequently showed the strongest spatial agreement with the intrinsic β-catenin-miRFP670 reference signal and the most favorable balance between detection coverage and specificity, whereas V5 remained useful as an orthogonal tag for immunostaining and biochemical detection.

Twin-prime editing further increases confidence in this fluorescent readout because the inserted sequence is distributed between two epegRNAs and reconstructed through coordinated editing. GFP11 complementation therefore indicates formation of a functional tagged product at the target locus, and because the S33Y substitution and GFP11 insertion are encoded within the same editing event, fluorescence reports the associated mutation with high confidence. GFP-positive cells could consequently be related directly to β-catenin redistribution, TOP reporter activation and tissue distortion. Thus, split-GFP provides a functional single-cell readout of productive editing without requiring clonal isolation.

Amplicon sequencing both supported this interpretation and highlighted the limitations of equating bulk read frequencies with cellular editing efficiency in a mosaic electroporation system. In the base- and prime-editing experiments, FACS enrichment relied on a co-electroporated H2B-RFP reporter rather than on the editing machinery itself. RFP-positive populations can therefore contain cells that received the reporter but not all independently delivered editing components, a concern that increases with the complexity of the editing system. WT reads may additionally originate from the unedited allele of successfully edited cells. Thus, the high WT fractions observed after RFP selection should not be interpreted directly as the proportion of cells in which editing failed. Instead, these experiments provide molecular confirmation of the intended alleles within populations whose apparent editing frequencies are influenced by mosaic plasmid delivery and allelic configuration.

This distinction was particularly informative for twin-prime editing. The exact S33Y–GFP11 allele increased from 0.04% of reads after H2B-RFP selection to 5.26% after GFP-based selection, representing more than 100-fold enrichment. When reads assigned to the edited reference but containing additional sequence variations were included, approximately 12% of reads in the GFP-selected population were associated with the S33Y–GFP11 reference. Moreover, the prominent deletions affecting the reconstructed region in the RFP-selected population were markedly reduced after GFP selection. GFP complementation therefore strongly enriches against incomplete editing products that disrupt functional GFP11 reconstruction, even though it does not yield a population composed exclusively of exact edited reads. This strong enrichment is consistent with the sequencing analysis described in the Results, where GFP-based isolation markedly increased recovery of the intended twinPE product.

Several factors can explain the remaining discrepancy between GFP positivity and the fraction of exact edited reads. GFP-positive cells can retain a WT second allele, consistent with the low frequency of GFP11/V5 double-positive cells in the competitive editing experiment, which indicates that independent editing of both alleles is infrequent. Exact-read classification is also stringent: sequencing errors or minor variations cause otherwise closely matching molecules to fall outside the exact S33Y– GFP11 category, an important consideration for Oxford Nanopore sequencing. In addition, the edited amplicon is longer than the WT amplicon (342 versus 264 bp), potentially favoring amplification of the shorter WT product during PCR, although the magnitude of such bias cannot be established here. Finally, the GFP-sorted population is unlikely to be absolutely pure. Doublet discrimination minimizes but cannot completely exclude cell aggregates, and the relatively weak fluorescence generated from endogenous GFP11 complementation limits separation between GFP-positive and negative populations. Despite conservative gating, a minor contribution from GFP-negative cells therefore cannot be excluded.

Importantly, functional GFP complementation itself does not require absolute nucleotide identity across the entire edited amplicon. Minor nucleotide changes that preserve an in-frame functional GFP11 peptide could remain fluorescent while being classified as sequence variants. GFP should therefore be regarded as a functional reporter of productive tag reconstruction rather than proof of nucleotide-perfect editing. Conversely, the imaging experiments provide information that sequencing cannot capture: GFP fluorescence reproduced the characteristic localization of β-catenin at adherens junctions and, for the S33Y allele, was accompanied by the expected cytoplasmic and nuclear accumulation, TOP reporter activation and epithelial phenotype. The combination of GFP complementation, appropriate subcellular localization and mutation-dependent functional effects therefore provides a high-confidence single-cell readout of productive endogenous editing, while sequencing remains necessary to characterize the full spectrum of molecular outcomes.

The competitive epegRNA experiment extended this approach to allelic analysis. Simultaneous delivery of GFP11- and V5-encoding epegRNA pairs allowed cells carrying different edits on the two β-catenin alleles to be identified. Although the assay cannot distinguish monoallelic editing from biallelic installation of the same tag, the low frequency of GFP11/V5 double-positive cells indicates that independent biallelic editing was infrequent. This is particularly relevant for modelling somatic tumor initiation, where an oncogenic mutation can arise in one allele of an individual cell surrounded by genetically normal neighbors. In this context, the mosaicism generated by in ovo electroporation becomes an experimental advantage rather than simply a limitation of delivery.

The endogenous tagging experiments also revealed an important design principle. GFP11 insertion within the β-TrCP phosphodegron partially stabilized β-catenin, whereas insertion immediately upstream at the WT-SP position retained predominantly junctional localization, did not induce detectable ectopic Wnt activation and preserved tissue architecture. Successful endogenous tagging must therefore be evaluated functionally rather than inferred from tag size or predicted structural tolerance. Even within intrinsically disordered regions, small insertions can perturb recognition motifs or protein turnover. The WT-SP site provides a validated reference position for comparing β-catenin variants under endogenous regulatory conditions.

This strategy should facilitate systematic analysis of disease-associated β-catenin mutations and may be applicable to other proteins whose function depends on dosage, localization or dynamic interactions. Introducing different mutations at the endogenous locus while retaining a common fluorescent tag should permit direct comparison of protein localization, signaling activity and effects on tissue organization without the confounding influence of heterogeneous overexpression.

Several limitations remain. Prime and twin-prime editing were less efficient than base editing, and their performance will depend on local sequence context, PAM availability, epegRNA architecture and delivery. Insert size also remains a practical constraint. Split-GFP complementation strongly enriches for productive editing but does not replace molecular characterization of genomic outcomes, particularly when nucleotide-level fidelity is itself the experimental endpoint. GFP1–10 must also be supplied exogenously, and variation in its expression may influence fluorescence intensity. Improvements in editor activity, guide design, delivery and sequencing should further increase the efficiency and generality of the approach.

Together, these results establish a framework in which precision editing can be used not only to modify an endogenous sequence but also to identify the edited cell, visualize the resulting protein and relate its behavior to signaling and tissue phenotype. In the chick neural tube, this enables analysis of the immediate consequences of somatic mutations without stable transgenesis, clonal selection or non-physiological protein expression. More broadly, coupling endogenous mutation to direct protein visualization provides a practical route to investigate how disease-associated alleles alter protein behavior within their native developmental and tissue context.

## METHODS

### Antibodies and chemicals

#### For immunohistochemistry

F-actin was labelled using Phalloidin Alexa Fluor Plus 647 (1:500, Thermo Fisher Scientific, Cat# A30107). Nuclei were counterstained with Hoechst 33342 (1:5,000, Thermo Fisher Scientific, Cat# H1399).

Primary antibodies: mouse anti-β-catenin clone 15B8 (1:500, Sigma-Aldrich, Cat# C7207, RRID_AB-476865), mouse anti-V5 antibody (1:500, Thermo Fisher Scientific, Cat# R961-25, RRID-AB-2556565).

Secondary antibodies: donkey anti-mouse Alexa Fluor 488 (Thermo Fisher Scientific Cat# A-21202, RRID AB-141607) goat anti-mouse Alexa Fluor 555 (Thermo Fisher Scientific, Cat# A-21422, RRID AB-2535844). All secondary antibodies were used at a dilution of 1:500.

#### For immunoblotting

Primary antibodies: rabbit monoclonal anti-β-catenin, clone M7A19 (1:500, Selleck, Cat# F2521, RRID: AB_3696821), mouse anti-V5 antibody (1:500, Thermo Fisher Scientific, Cat# R961-25, RRID AB-2556565).

Secondary antibodies: goat anti-mouse DyLight-800 (Thermo Fisher scientific Cat# SA5-35521, RRID AB-2556774), goat anti-rabbit DyLight-680 (Thermo Fisher Scientific Cat# 35568 RRID AB-614946)

### DNA constructs

To generate sgRNAs for base editing, forward (Fw) and reverse (Rv) oligonucleotides were annealed and ligated into the BbsI–BbsI cloning site of pGuia. To generate epegRNAs for prime editing or twin-prime editing, the spacer, scaffold, and PBS–RT regions were ordered separately. Forward and reverse oligonucleotides were annealed and assembled by Golden Gate (GG)-based multi-fragment ligation into the BsaI–BsaI cloning site of the pU6-tevopreQ1 vector. To generate the pTest 1×ALFA, 1×V5, and 1×GFP11 constructs, the tag sequences were ordered as forward and reverse oligonucleotides, annealed, and ligated into the BsmBI–BsmBI cloning site of pTest. To generate the pTest 3×ALFA, 3×V5, 2×GFP11 and 3×GFP11 constructs, the tag sequences were ordered as double-stranded DNA fragments (GeneArt™ Strings™, Thermo Fisher Scientific) and ligated into the BsmBI–BsmBI cloning site of pTest. All remaining constructs were generated using conventional restriction enzyme digestion and ligation methods. The parental vectors used in this study are listed below. Detailed names and sequences of all derived constructs, oligonucleotides, and DNA inserts are provided in Supplementary Table 1.

#### Base Editing

pCMV-CBE6b-SpCas9-D10A-2xUGI (CBE6b): sixth-generation cytidine base editor, Liu Lab. Addgene #215820.

pGuia: U6-driven sgRNA expression vector carrying an eGFP reporter, this work.

#### Prime Editing

pCMV-PEmax (PEmax): SpCas9(H840A)-MMLV reverse transcriptase prime editor, Liu Lab. Addgene #174820.

pU6-tevopreQ1-GG-Acceptor (pU6-TevQ1): epegRNA expression vector, Liu Lab. Addgene #174038.

pbGuia: U6-driven sgRNA expression vector lacking the eGFP reporter, this work.

#### Twin Prime Editing

pCMV-PEmax (PEmax): SpCas9(H840A)-MMLV reverse transcriptase prime editor, Liu Lab. Addgene #174820.

pU6-tevopreQ1-GG-Acceptor (pU6-TevQ1): epegRNA expression vector, Liu Lab. Addgene #174038.

#### Tag Selection

pTest: pCIEGO-derived β-catenin S33Y–miRFP670 reporter containing a BsmBI–BsmBI Golden Gate cloning cassette between residues Y33 and G34 for tag insertion, this work.

#### Nanobodies

pmCherryN1-NbV5: expresses NbV5–mCherry, Giguère Lab. Addgene #201474. pEGFPN1-NbV5: expresses NbV5–eGFP, Giguère Lab. Addgene #201473.

pCDNA6/my-His-NbALFA: expresses NbALFA–GFP2, Giguère Lab. Addgene #201485.

pCDNA3.1-NbALFA: expresses GFP2–NbALFA, Giguère Lab. Addgene #201486. pmiRFP670nano3N1-NbV5: expresses NbV5–miRFP670nano3, this work.

#### Split GFP

pHR-SFFV-GFP1-10: expresses GFP1–10, Huang Lab. Addgene #40809.

pCIEGO-GFP1-10: expresses GFP1–10, this work.

#### Other Vectors

pCIG: CMV-IE enhancer/chicken β-actin promoter IRES-GFP expression vector (Megason and McMahon, 2002).

pCIEGO: CMV-IE enhancer/chicken β-actin promoter expression vector (Herrera et al., 2014).

TOP-Flash: 5× TCF/LEF-responsive elements upstream of a TK minimal promoter driving luciferase expression (Korinek et al., 1997).

TOP-RFP: 5× TCF/LEF-responsive elements upstream of a TK minimal promoter driving H2B-RFP expression (Herrera et al., 2023).

pSHIN: H1-driven shRNA expression vector carrying an eGFP reporter (Kojima and Borisy, 2014).

pCS2+-mbGFP: expresses membrane-targeted GFP (Das and Storey, 2012).

pCMV-PACT-RFP: expresses the PACT domain fused to RFP as a centriolar marker (Gillingham and Munro, 2000).

### FACS isolation and amplicon sequencing of genome-editing outcomes

To characterize genome-editing outcomes at the endogenous β-catenin locus, HH12 chick embryos were electroporated with the corresponding editing components together with pCAGGS-IRES-H2B-RFP as a marker of transfected cells. Three editing strategies were analysed: base editing to introduce the S33F substitution, prime editing to introduce S33Y, and twin-prime editing to introduce S33Y together with a GFP11 tag flanked by 5×G linkers between residues L31 and D32 of β-catenin. Fifteen embryos were pooled for base editing, 20 for prime editing, 17 for RFP-selected twin-prime editing, and 23 for GFP-selected twin-prime editing.

At 24 h after electroporation, the electroporated region of the neural tube was dissected in cold PBS and pooled for each experimental condition. Tissues were dissociated in 0.25% trypsin-EDTA (Gibco, Cat# 25200-07) supplemented with DNase I (Sigma, Cat# D5025; approximately 100 μg/mL final concentration). Samples were incubated for 10 min on ice followed by 15 min at 37 °C with agitation at 850 rpm. Digestion was stopped on ice by addition of DMEM containing 20% FBS, and tissues were mechanically dissociated by pipetting. Undissociated tissue fragments were allowed to settle before transferring the cell suspension to a fresh tube.

Cells were sorted using a BD FACS Fusion II cell sorter. For base editing, prime editing and twin-prime editing, H2B-RFP-positive cells were isolated to enrich for electroporated cells, and 50,000 cells were collected for each condition. For twin-prime editing, an additional population was sorted on the basis of GFP fluorescence generated by GFP11/GFP1–10 complementation, and approximately 60,000 GFP-positive cells were collected.

Sorted cells were centrifuged at 800 × g for 5 min at 4 °C and genomic DNA was extracted in 50 μL of buffer containing 10 mM Tris-HCl (pH 8.0), 0.1 mM EDTA, 0.5% Tween-20 and 100 μg/mL Proteinase K (Thermo Fisher Scientific, Cat# E00491). Samples were incubated at 56 °C for 16 h, followed by 10 min at 95 °C to inactivate Proteinase K.

The genomic region encompassing exon 4 of chicken β-catenin was amplified in duplicate using Q5 Hot Start High-Fidelity DNA Polymerase (NEB, Cat# M0493) and primers located in the flanking intronic sequences (forward, 5′-TTCAACGATTTCTTACAG-3′; reverse, 5′-TTGTAGAGAAGGCCTTAC-3′). The resulting amplicon was 264 bp for the WT and single-nucleotide-edited alleles and 342 bp for the S33Y-GFP11 allele. PCR conditions were 98 °C for 30 s, followed by 35 cycles of 98 °C for 10 s, 60 °C for 30 s and 72 °C for 20 s, with a final extension at 72 °C for 10 min. PCR products were submitted to Plasmidsaurus for Standard Premium PCR sequencing with cleanup using the Oxford Nanopore platform.

Raw FASTQ files were analysed using online version of CRISPResso2 (version 2.3.4; https://crispresso2.pinellolab.org). For base-editing analysis, reads were aligned to the WT β-catenin exon 4 amplicon using the sgRNA sequence CTGGACTCTGGTATCCATTC. CRISPResso2 was configured for base-editor analysis (C-to-T conversion), using a 30-bp quantification window centered 10 bp upstream of the predicted nCas9(D10A) cleavage site and a 30-bp plotting window. For prime editing, reads were aligned to the WT amplicon using the spacer sequence CAGCAGTCATATCTGGACTC, with a 30-bp quantification window centered 3 bp upstream of the predicted nCas9(H840A) cleavage site and a 30-bp plotting window. For twin-prime editing, reads were aligned against both the WT and expected S33Y-GFP11 amplicon sequences using the guide sequence TTGGCAGCAGCAGTCATATC. A 67-bp quantification window centered 47 bp downstream of the predicted cleavage site and a 67-bp plotting window were used for both H2B-RFP- and GFP-selected samples. Other parameters were left at their default settings.

### Chick embryo *in ovo* electroporation

Fertilized White Leghorn chicken eggs were incubated at 38.5 °C in 45% humidity. Embryos were staged according to the Hamburger and Hamilton (HH) staging system (Hamburger and Hamilton, 1992). Column-purified plasmid DNA was diluted to the concentrations indicated for each experiment in sterile H₂O containing 0.05% Fast Green. DNA was injected into the lumen of the neural tube of HH12 embryos, and electroporation was performed by applying five 50-ms square-wave pulses at 25 V using an Intracel TSS20 electroporator with electrodes positioned on either side of the neural tube.

Following electroporation, embryos were returned to the incubator until the indicated developmental stage and subsequently dissected under a fluorescence stereomicroscope. Under these conditions, embryos electroporated at HH12 typically reached stages HH18 and HH23 after 24 and 48 h, respectively. Embryos that failed to reach the expected developmental stage were excluded from further analysis.

### Immunostaining

Embryos were fixed overnight at 4 °C in 4% paraformaldehyde (PFA) in phosphate-buffered saline (PBS). Samples were either sectioned at a thickness of 60 μm using a VT1000S vibratome (Leica) or opened longitudinally along the roof plate to generate open-book preparations.

Immunostaining was performed using standard procedures. Samples were washed in PBT (PBS containing 0.1% Triton X-100), incubated with the appropriate primary antibodies and subsequently labelled with Alexa Fluor- or cyanine-conjugated secondary antibodies. Stained sections and open-book preparations were mounted in Fluoromount (Sigma-Aldrich).

Images were acquired at 18 °C using a Dragonfly 500 spinning-disk confocal microscope (Oxford Instruments) equipped with 40×, 60× or 100× objectives. Image processing and analysis were performed using Fiji/ImageJ.

### Immunoblotting

Electroporated neural tubes were dissected from chicken embryos and lysed in PIK buffer (20 mM Tris-HCl, pH 7.4, 137 mM NaCl, 10% glycerol, 1% NP-40, 1 mM CaCl₂ and 1 mM MgCl₂) supplemented with protease inhibitors (1 mM PMSF, 10 μg/mL aprotinin and 10 μg/mL leupeptin). Insoluble material was removed by centrifugation, and the supernatants were mixed with 5× Laemmli sample buffer to a final 1× concentration (65 mM Tris-HCl, pH 6.8, 2% SDS, 10% glycerol, 100 mM DTT and 0.5 mg/mL bromophenol blue). Samples were heated at 95 °C for 5 min, resolved by SDS-PAGE on 8% polyacrylamide gels and transferred onto Immobilon-FL PVDF membranes (Millipore).

Membranes were blocked with Odyssey Blocking Buffer (TBS) (LI-COR Biosciences, Cat. No. 927-50000) and incubated with primary antibodies diluted in the same buffer supplemented with 0.2% Tween-20. Following three washes in TTBS (20 mM Tris-HCl, pH 7.4, 150 mM NaCl and 0.1% Tween-20), membranes were incubated with IRDye® 680RD- or IRDye® 800CW-conjugated secondary antibodies diluted in Odyssey Blocking Buffer containing 0.2% Tween-20 and 0.01% SDS. After three additional washes in TTBS, membranes were air-dried and scanned using an Odyssey Infrared Imaging System (LI-COR Biosciences). Molecular weights were estimated using Precision Plus Protein™ Standards (Bio-Rad)

### *In vivo* luciferase reporter assay

Embryos were electroporated with the indicated DNA constructs together with the 5×TCF-binding site luciferase reporter (TOPFlash) containing synthetic TCF-binding sites (Korinek et al., 1997), and a Renilla luciferase expression plasmid (Promega) as an internal normalization control. At 48 h post-electroporation (hpe), GFP-positive neural tubes were dissected and homogenized in Passive Lysis Buffer. Firefly and Renilla luciferase activities were measured using the Dual-Luciferase® Reporter Assay System (Promega, Cat. No. E1910).

### Quantitative evaluation of GFP11, ALFA and V5 tag performance

To quantitatively compare the performance of the ALFA, V5 and GFP11 tags, confocal images were acquired 24 h after electroporation and analysed in Fiji. Six independent embryos were analysed per condition, using one confocal image per embryo. For each image, a maximum-intensity projection of five Z-planes encompassing the adherens junction level was generated. A single 25 × 25 μm region of interest (ROI), containing a group of predominantly transfected cells, was selected from each projection and used for all subsequent quantitative analyses.

Background fluorescence was removed using the Sliding Paraboloid algorithm (radius = 50 pixels). Colocalization between the intrinsic β-catenin–miRFP670 signal and the corresponding tag-dependent fluorescence was then quantified within each ROI using the Coloc2 plugin.

Pearson’s correlation coefficient was used to assess the spatial similarity between the two signals. Thresholded Manders’ coefficients (tM1 and tM2) were used to quantify the fraction of β-catenin signal detected by each tag and the fraction of tag-dependent fluorescence corresponding to bona fide β-catenin localization, respectively. Thresholds were determined automatically using the Costes randomization method.

In addition, the regression slope obtained from pixel-by-pixel intensity correlation analysis was used to assess the proportionality between β-catenin–miRFP670 and tag-dependent fluorescence intensities. Together, these parameters provided a quantitative assessment of tag fidelity, detection coverage and signal specificity.

### Quantification of twin-prime editing efficiency of b-CatS33Y-GFP11 Edit-1 and Edit-2

Twin-prime editing efficiency was quantified from confocal images acquired 24 h after electroporation. Six independent embryos were analysed per condition, generating 15 fields of 104.6 × 104.8 μm for each editing strategy. For each field, a maximum-intensity projection of five Z-planes encompassing the centrosomal layer was generated and analysed in Fiji.

To determine the total number of transfected cells, the PACT-RFP channel was subjected to background subtraction using the Sliding Paraboloid algorithm (radius = 50 pixels), and PACT-RFP-positive centrosomes were automatically detected using the TrackMate plugin (expected spot size = 0.5 μm; quality threshold = 1). GFP11-positive edited cells were counted manually in the corresponding GFP channel using the Cell Counter plugin. Editing efficiency was calculated independently for each field as the percentage of GFP11-positive cells relative to the total number of PACT-RFP-positive cells.

### Quantification of GFP11/V5 competitive editing

Competitive twin-prime editing outcomes were quantified from confocal images acquired 24 h after electroporation. Six independent embryos were analysed, generating a total of 16 fields of 104.6 × 104.8 μm. For each field, a maximum-intensity projection of five Z-planes encompassing the adherens junction level was generated and analysed in Fiji. GFP11-positive, V5-positive and GFP11/V5 double-positive cells were counted manually using the Cell Counter plugin. For each field, the frequencies of GFP11- and V5-positive cells were calculated relative to the combined number of GFP11- and V5-positive cells, whereas GFP11/V5 double-positive cells were quantified separately relative to the same denominator.

### Quantification of junctional and nuclear GFP11-labelled β-catenin

Four independent embryos were analysed per condition, with one confocal stack acquired per embryo. Quantification of edited β-catenin fluorescence was performed in Fiji using confocal stacks comprising 33 optical sections spanning 4.95 μm (field size: 104.6 × 104.8 μm).

For quantification of junctional β-catenin, maximum-intensity projections of five consecutive Z-planes centered on the apical adherens junction (AJ) layer were generated. Ten 20 × 20 μm subregions centered on clusters of transfected cells were selected from each image. Within each subregion, the GFP channel corresponding to GFP11-tagged endogenous β-catenin revealed by GFP1–10 complementation was thresholded to generate a binary mask encompassing the adherens junctions. The resulting mask was converted into a region of interest (ROI), applied to the original image, and the mean gray value of the GFP signal was measured within the masked junctional area. A total of 40 AJ measurements were obtained per condition.

For quantification of nuclear β-catenin, maximum-intensity projections of five consecutive Z-planes centered on the nuclear layer were generated from the same confocal stacks. Ten circular ROIs (2.5 μm diameter) were manually placed on GFP-positive nuclei in each image using DAPI staining to identify nuclear boundaries. The mean gray value of the GFP signal was measured within each ROI, yielding a total of 40 nuclear measurements per condition.

For statistical analysis, the ten measurements obtained from each embryo were averaged, and embryo means (*n* = 4 per condition) were compared separately for AJ and nuclear fluorescence using one-way ANOVA followed by Dunnett’s multiple-comparisons test.

### Statistics

All statistical analyses were performed using GraphPad Prism 6.

For **Fig. 1N**, luciferase activity following β-catenin S33Y misexpression was analysed using an unpaired two-tailed Student’s *t*-test, whereas base-editing conditions were compared using one-way ANOVA followed by Dunnett’s multiple-comparisons test.

For **Fig. 2J**, luciferase activity was analysed using one-way ANOVA followed by Tukey’s multiple-comparisons test.

For **Fig. 4K**, the percentage of correctly edited cells among transfected cells was analysed using an unpaired two-tailed Student’s *t*-test.

For **Fig. 6P**, mean GFP fluorescence intensity at adherens junctions and in nuclei was analysed by one-way ANOVA followed by Tukey’s multiple-comparisons test and Dunnett’s multiple-comparisons test, respectively.

For **Fig. 6Y**, luciferase activity following base editing was analysed using an unpaired two-tailed Student’s *t*-test, whereas prime- and twin-prime-editing conditions were analysed using one-way ANOVA followed by Dunnett’s multiple-comparisons test.

Unless otherwise indicated, data are presented as mean ± SEM. Statistical significance was defined as *P* < 0.05

## Supporting information

Supplementary Figures and Tables with legends

## ACKNOWLEDGEMENTS

We thank Patrick Giguère (University of Ottawa) for kindly providing the GFP2-NbALFA (Addgene plasmid #201486), NbALFA-GFP2 (#201485), NbV5-eGFP (#201473) and NbV5-mCherry (#201474) constructs, and Bo Huang for kindly providing the GFP1–10 expression plasmid (Addgene plasmid #40809).

This work was supported by grant PID2023-146672NB-I00 funded by MICIU/AEI/10.13039/501100011033.

## AUTHOR CONTRIBUTIONS

A.M. and C.R. contributed equally to this work. S.P. conceived the study and designed the overall experimental strategy. A.M. and C.R. contributed equally to experimental design and performed the experimental work, including molecular cloning and construct design and validation, DNA preparation, in ovo electroporation and embryo manipulation, immunohistochemistry, image acquisition, luciferase assays and western blotting. S.P. performed image analysis, statistical analysis and data visualization, and prepared the figures. A.M., C.R. and S.P. interpreted and discussed the results. S.P. supervised the study, acquired funding and wrote the original manuscript. A.M. and C.R. reviewed and edited the manuscript.

