## Supplementary Figures and Tables with legends for "Twin-prime editing enables endogenous protein visualization and mutant allele tracking"

Supp Fig1

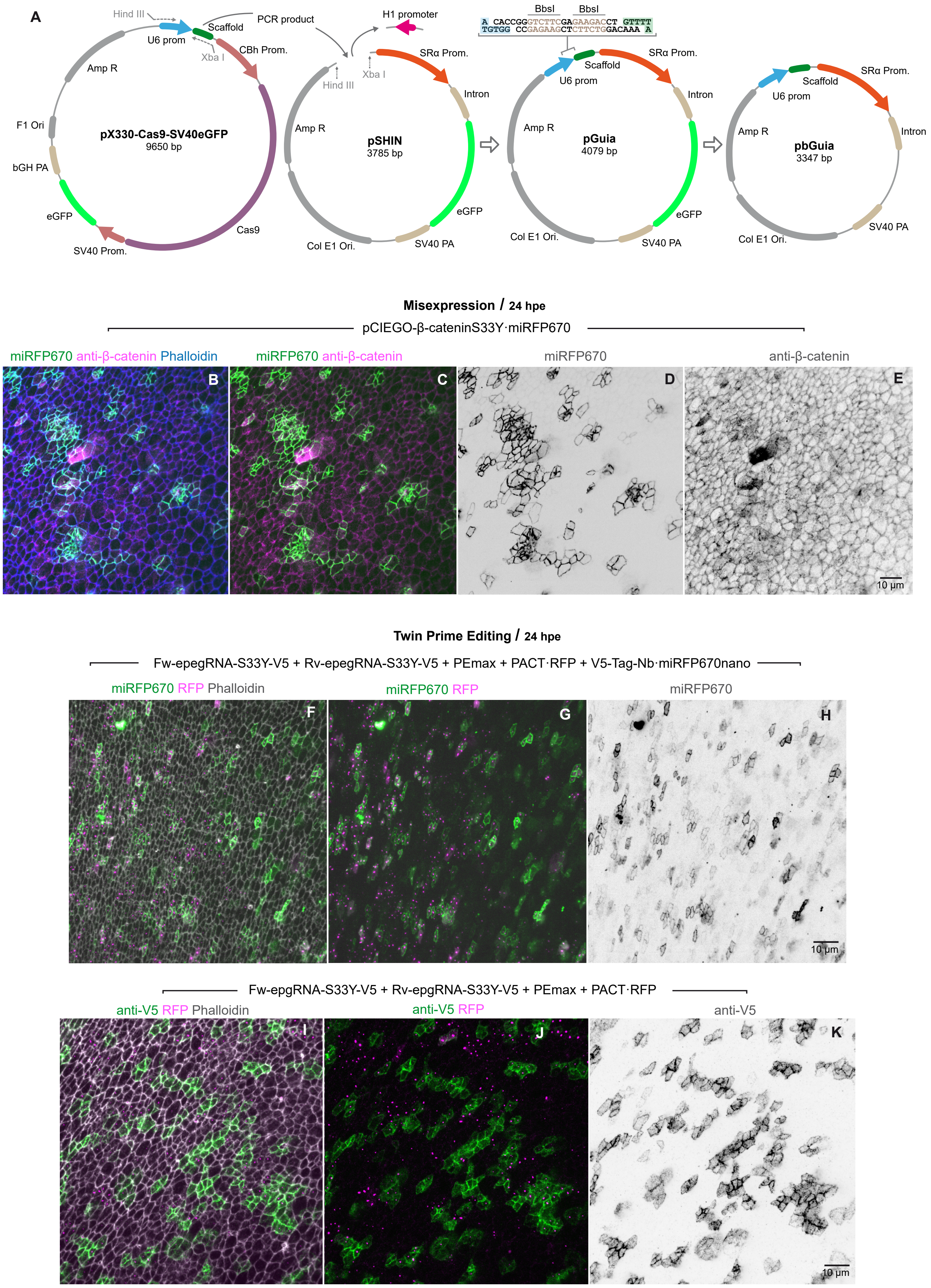
A
Hind III
U6 prom
Scaffold
Xba I
PCR product
CBh Prom.
Amp R
F1 Ori
pX330-Cas9-SV40eGFP
9650 bp
bGH PA
eGFP
SV40 Prom.
Cas9
H1 promoter
Hind III
Xba I
SRα Prom.
Intron
Amp R
pSHIN
3785 bp
eGFP
Col E1 Ori.
SV40 PA
BbsI
BbsI
A CACCGGGTCTTCGAGAAGACCT GTTTT
TGTGG CCGAGAAGCTCTTCTGGACAAA A
Scaffold
U6 prom
SRα Prom.
Intron
Amp R
pGuia
4079 bp
eGFP
Col E1 Ori.
SV40 PA
Scaffold
U6 prom
SRα Prom.
Intron
Amp R
pbGuia
3347 bp
Col E1 Ori.
SV40 PA
Misexpression / 24 hpe
pCIEGO-β-cateninS33Y·miRFP670
miRFP670 anti-β-catenin Phalloidin
miRFP670 anti-β-catenin
miRFP670
anti-β-catenin
B
C
D
E
10 µm
Twin Prime Editing / 24 hpe
Fw-epegRNA-S33Y-V5 + Rv-epegRNA-S33Y-V5 + PEmax + PACT·RFP + V5-Tag-Nb·miRFP670nano
miRFP670 RFP Phalloidin
miRFP670 RFP
miRFP670
F
G
H
10 µm
Fw-epgRNA-S33Y-V5 + Rv-epgRNA-S33Y-V5 + PEmax + PACT·RFP
anti-V5 RFP Phalloidin
anti-V5 RFP
anti-V5
I
J
K
10 µm

Supp Fig2

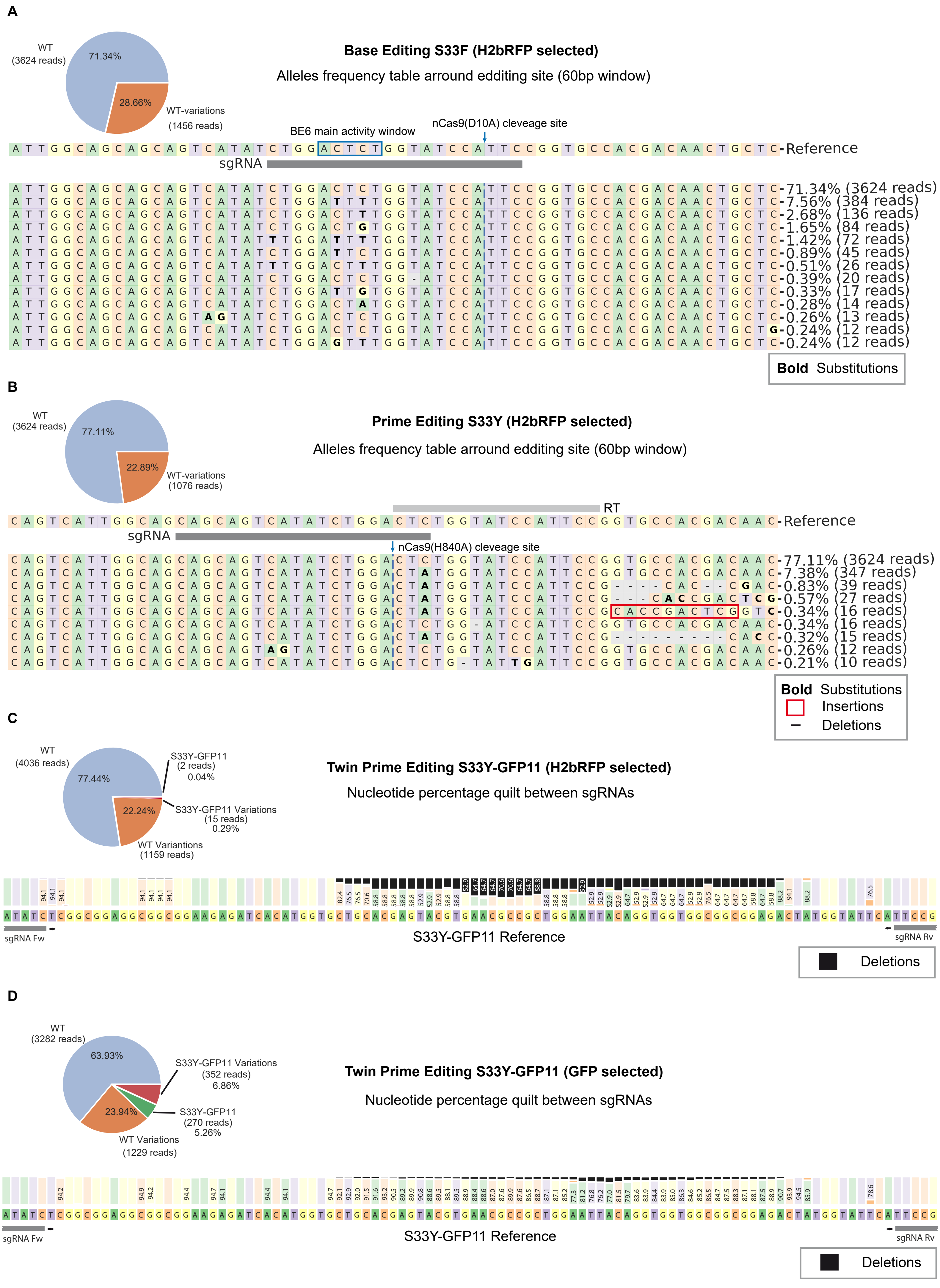
A
Base Editing S33F (H2bRFP selected)
Alleles frequency table arround eddíting site (60bp window)
WT
(3624 reads)
71.34%
28.66%
WT-variations
(1456 reads)
BE6 main activity window
nCas9(D10A) cleveage site
Reference
sgRNA
71.34% (3624 reads)
7.56% (384 reads)
2.68% (136 reads)
1.65% (84 reads)
1.42% (72 reads)
0.89% (45 reads)
0.51% (26 reads)
0.39% (20 reads)
0.33% (17 reads)
0.28% (14 reads)
0.26% (13 reads)
0.24% (12 reads)
0.24% (12 reads)
Bold Substitutions
B
Prime Editing S33Y (H2bRFP selected)
Alleles frequency table arround eddíting site (60bp window)
WT
(3624 reads)
77.11%
22.89%
WT-variations
(1076 reads)
RT
Reference
sgRNA
nCas9(H840A) cleveage site
77.11% (3624 reads)
7.38% (347 reads)
0.83% (39 reads)
0.57% (27 reads)
0.34% (16 reads)
0.34% (16 reads)
0.32% (15 reads)
0.26% (12 reads)
0.21% (10 reads)
Bold Substitutions
Insertions
Deletions
C
Twin Prime Editing S33Y-GFP11 (H2bRFP selected)
Nucleotide percentage quilt between sgRNAs
WT
(4036 reads)
77.44%
22.24%
S33Y-GFP11
(2 reads)
0.04%
S33Y-GFP11 Variations
(15 reads)
0.29%
WT Variantions
(1159 reads)
sgRNA Fw
S33Y-GFP11 Reference
sgRNA Rv
Deletions
D
Twin Prime Editing S33Y-GFP11 (GFP selected)
Nucleotide percentage quilt between sgRNAs
WT
(3282 reads)
63.93%
23.94%
S33Y-GFP11 Variations
(352 reads)
6.86%
S33Y-GFP11
(270 reads)
5.26%
WT Variations
(1229 reads)
sgRNA Fw
S33Y-GFP11 Reference
sgRNA Rv
Deletions

**Supplementary Figure 1. Generation of pGuia and comparison of β-catenin detection following misexpression and V5 twin-prime editing.**

**(A)** Schematic representation of the generation of the pGuia sgRNA-expression vector. The U6 promoter and sgRNA scaffold from pX330-Cas9-SV40eGFP were amplified by PCR and inserted into the pSHIN backbone to replace the H1 promoter, generating pGuia. A GFP-negative derivative (pbGuia) was generated by removal of the GFP expression cassette.

**(B–E)** Open-book preparation of chick neural tube electroporated at HH12 with pCIEGO-β-catenin-S33Y-miRFP670 and analysed 24 hpe. (B) Merge showing intrinsic miRFP670 fluorescence (green), anti-β-catenin immunostaining (magenta) and phalloidin (blue). (C) Merge of miRFP670 fluorescence and anti-β-catenin immunostaining. (D) miRFP670 channel. (E) Anti-β-catenin immunostaining. The intrinsic miRFP670 signal reproduces the distribution detected by the β-catenin antibody.

**(F–H)** Detection of endogenous β-catenin edited by twin-prime editing using the V5 chromobody. Embryos were electroporated at HH12 with PEmax, the S33Y-V5 forward and reverse epegRNAs, PACT-RFP and the V5-Tag nanobody fused to miRFP670nano and analysed 24 hpe. (F) Merge showing V5 chromobody fluorescence (green), PACT-RFP (magenta) and phalloidin (grey). (G) Merge of V5 chromobody fluorescence and PACT-RFP. (H) V5 chromobody fluorescence.

**(I–L)** Detection of endogenous β-catenin edited by twin-prime editing using anti-V5 immunofluorescence. Embryos were electroporated at HH12 with PEmax, the S33Y-V5 forward and reverse epegRNAs and PACT-RFP and analysed 24 hpe. (I) Merge showing anti-V5 immunostaining (green), PACT-RFP (magenta) and phalloidin (grey). (J) Merge of anti-V5 immunostaining and PACT-RFP. (K) Anti-V5 immunostaining. (L) Schematic comparison of V5 detection by fluorescent chromobody or immunostaining following twin-prime editing.

**Supplementary Figure 2. Amplicon sequencing analysis of endogenous β-catenin editing outcomes.**

**(A)** CRISPResso2 analysis of the β-catenin exon 4 amplicon following S33F base editing in H2B-RFP-positive cells isolated by FACS 24 hpe. The pie chart shows the proportion of reads corresponding to the WT sequence or containing sequence variations. The allele-frequency table shows the most abundant sequences within a 60-bp window surrounding the editing site. The sgRNA sequence, BE6 main activity window and predicted nCas9(D10A) cleavage site are indicated.

**(B)** CRISPResso2 analysis of S33Y prime editing in H2B-RFP-positive cells isolated by FACS 24 hpe. The pie chart shows the proportion of WT and sequence-variant reads, and the allele-frequency table shows the most abundant sequences within a 60-bp window surrounding the editing site. The spacer sequence, reverse-transcription (RT) region and predicted nCas9(H840A) cleavage site are indicated.

**(C)** CRISPResso2 analysis of S33Y-GFP11 twin-prime editing in H2B-RFP-positive cells isolated by FACS 24 hpe. Reads were aligned against both the WT and expected S33Y-GFP11 reference sequences. The pie chart shows reads classified as WT, WT containing sequence variations, S33Y-GFP11, or S33Y-GFP11 containing additional sequence variations. The nucleotide-percentage quilt shows sequence composition across the region between the two epegRNA target sites relative to the expected S33Y-GFP11 reference sequence.

**(D)** CRISPResso2 analysis of S33Y-GFP11 twin-prime editing following FACS enrichment (24 hpe) based on GFP fluorescence generated by GFP11/GFP1–10 complementation. Reads were analysed as in **C**. The pie chart shows the distribution of reads assigned to the WT and S33Y-GFP11 reference sequences, with or without additional sequence variations, and the nucleotide-percentage quilt shows sequence composition across the region between the two epegRNA target sites relative to the expected S33Y-GFP11 reference sequence.

### Supplementary Table 1. Oligonucleotides and synthetic DNA fragments used in this study.

| | Construct number | Construct name | Oligo number | Oligo name | Direction | Sequence (5'-3') |
|---|---|---|---|---|---|---|
| Base Editing | 1205 | pGUÍA-Scrb | 107 | U6-prom-HindIII | Fw | ATAAAGCTTGAGGGCCTATTTCCCATGAT |
| | | | 108 | U6-prom-XbaI | Rv | GGTACCTCTAGAGCCATTTG |
| | 1210 | pGUÍA β-Catenina S33F | 109 | gCTNNB1 FW | Fw | CACCGCTGGACTCTGGTATCCATTC |
| | | | 110 | gCTNNB1 RV | Rv | AAACGAATGGATACCAGAGTCCAGC |
| Prime Editing | 1247 | pU6-TevQ1-β-Cat-S33Y(+3) | 124 | Spacer β-Cat S33Y +3 | Fw | CACCGCAGCAGTCATATCTGGACTCGTTTT |
| | | | 125 | Spacer β-Cat S33Y +3 | Rv | CTCTAAAACGAGTCCAGATATGACTGCTGC |
| | | | 126 | scaffold | Fw | AGAGCTAGAAATAGCAAGTTAAAATAAGGCTAGTCCGTTATCAACTTGAAAAAGTGGGACCGAGTCG |
| | | | 127 | scaffold | Rv | GCACCGACTCGGTCCCACTTTTTCAAGTTGATAACGGACTAGCCTTATTTTAACTTGCTATTTCTAG |
| | | | 128 | PBS-RT-HA S33Y | Fw | GTGCGGAATGGATACCATAGTCCAGATATGACT |
| | | | 129 | PBS-RT-HA S33Y | Rv | CGCGAGTCATATCTGGACTATGGTATCCATTCC |
| | 1249 | pGUÍA 2º Cut | 130 | 2º Cut S33Y/S37Y | Fw | CACCGGTGTCCACATCTTCCTCTTC |
| | | | 131 | 2º Cut S33Y/S37Y | Rv | AAACGAAGAGGAAGATGTGGACACC |
| Twin Prime Editing | 1277 | pU6-TevQ1-β-Cat-S33Y-GFP11-Edit Fw | 124 | F Spacer β-Cat S33Y +3 | Fw | CACCGCAGCAGTCATATCTGGACTCGTTTT |
| | | | 125 | F Spacer β-Cat S33Y +3 | Rv | CTCTAAAACGAGTCCAGATATGACTGCTGC |
| | | | 126 | scaffold | Fw | AGAGCTAGAAATAGCAAGTTAAAATAAGGCTAGTCCGTTATCAACTTGAAAAAGTGGGACCGAGTCG |
| | | | 127 | scaffold | Rv | GCACCGACTCGGTCCCACTTTTTCAAGTTGATAACGGACTAGCCTTATTTTAACTTGCTATTTCTAG |
| | | | 181 | F PBS-RT-GFP11 S33Y | Fw | GTGCATTCCAGCGGCGTTCACGTACTCGTGCAGCACCATGTGATCTCTTCCGCCGCCTCCGCCATAATCCAGATATGACT |
| | | | 182 | F PBS-RT-GFP11 S33Y | Rv | CGCGAGTCATATCTGGATTATGGCGGAGGCGGCGGAAGAGATCACATGGTGCTGCACGAGTACGTGAACGCCGCTGGAAT |
| | 1278 | pU6-TevQ1-β-Cat-S33Y-GFP11-Edit-1 | 179 | R Spacer β-Cat S33Y | Fw | CACCGGGGAGCAGTTGTCGTGGCACGTTTT |
| | | | 180 | R Spacer β-Cat S33Y | Rv | CTCTAAAACGTGCCACGACAACTGCTCCCC |
| | | | 126 | scaffold | Fw | AGAGCTAGAAATAGCAAGTTAAAATAAGGCTAGTCCGTTATCAACTTGAAAAAGTGGGACCGAGTCG |
| | | | 127 | scaffold | Rv | GCACCGACTCGGTCCCACTTTTTCAAGTTGATAACGGACTAGCCTTATTTTAACTTGCTATTTCTAG |
| | | | 183 | R1 PBS-RT-GFP11 S33Y | Fw | GTGCGTACGTGAACGCCGCTGGAATTACAGGTGGTGGCGGCGGAGGTATCCATTCCGGAGCCACGACAACTGC |
| | | | 184 | R1 PBS-RT-GFP11 S33Y | Rv | CGCGGCAGTTGTCGTGGCTCCGGAATGGATACCTCCGCCGCCACCACCTGTAATTCCAGCGGCGTTCACGTAC |
| | 1279 | pU6-TevQ1-β-Cat-S33Y-GFP11-Edit-2 | 160 | Spacer β-Cat T41A+1 | Fw | CACCGCAAAGAGGGAGCAGTTGTCGGTTTT |
| | | | 161 | Spacer β-Cat T41A+1 | Rv | CTCTAAAACCGACAACTGCTCCCTCTTTGC |
| | | | 126 | scaffold | Fw | AGAGCTAGAAATAGCAAGTTAAAATAAGGCTAGTCCGTTATCAACTTGAAAAAGTGGGACCGAGTCG |
| | | | 127 | scaffold | Rv | GCACCGACTCGGTCCCACTTTTTCAAGTTGATAACGGACTAGCCTTATTTTAACTTGCTATTTCTAG |
| | | | 185 | R2 Spacer β-Cat S33Y | Rv | GTGCGTACGTGAACGCCGCTGGAATTACAGGTGGTGGCGGCGGAGGTATCCATTCCGGTGCAACTACAACTGCTCCCTC |
| | | | 186 | R2 Spacer β-Cat S33Y | Fw | CGCGGAGGGAGCAGTTGTAGTTGCACCGGAATGGATACCTCCGCCGCCACCACCTGTAATTCCAGCGGCGTTCACGTAC |
| | 1280 | pU6-TevQ1-β-Cat-S33Y-V5-Edit Fw | 124 | F Spacer β-Cat S33Y +3 | Fw | CACCGCAGCAGTCATATCTGGACTCGTTTT |
| | | | 125 | F Spacer β-Cat S33Y +3 | Rv | CTCTAAAACGAGTCCAGATATGACTGCTGC |
| | | | 126 | scaffold | Fw | AGAGCTAGAAATAGCAAGTTAAAATAAGGCTAGTCCGTTATCAACTTGAAAAAGTGGGACCGAGTCG |
| | | | 127 | scaffold | Rv | GCACCGACTCGGTCCCACTTTTTCAAGTTGATAACGGACTAGCCTTATTTTAACTTGCTATTTCTAG |
| | | | 187 | F PBS-RT-S33Y V5 | Fw | GTGCGCTATCCAGGCCGAGCAGTGGATTTGGAATGGGCTTGCCTCCGCCGCCTCCGCCATAATCCAGATATGACT |
| | | | 188 | F PBS-RT-S33Y V5 | Rv | CGCGAGTCATATCTGGATTATGGCGGAGGCGGCGGAGGCAAGCCCATTCCAAATCCACTGCTCGGCCTGGATAGC |
| | 1281 | pU6-TevQ1-β-Cat-S33Y-V5-Edit-1 | 179 | R Spacer β-Cat S33Y | Fw | CACCGGGGAGCAGTTGTCGTGGCACGTTTT |
| | | | 180 | R Spacer β-Cat S33Y | Rv | CTCTAAAACGTGCCACGACAACTGCTCCCC |
| | | | 126 | scaffold | Fw | AGAGCTAGAAATAGCAAGTTAAAATAAGGCTAGTCCGTTATCAACTTGAAAAAGTGGGACCGAGTCG |
| | | | 127 | scaffold | Rv | GCACCGACTCGGTCCCACTTTTTCAAGTTGATAACGGACTAGCCTTATTTTAACTTGCTATTTCTAG |
| | | | 189 | R1 PBS-RT-V5 | Fw | GTGCTCCACTGCTCGGCCTGGATAGCACAGGTGGTGGCGGCGGAGGTATCCATTCCGGAGCCACGACAACTGC |
| | | | 190 | R1 PBS-RT-V5 | Rv | CGCGGCAGTTGTCGTGGCTCCGGAATGGATACCTCCGCCGCCACCACCTGTGCTATCCAGGCCGAGCAGTGA |
| | 1282 | pU6-TevQ1-β-Cat-S33Y-V5-Edit-2 | 160 | Spacer β-Cat T41A+1 | Fw | CACCGCAAAGAGGGAGCAGTTGTCGGTTTT |
| | | | 161 | Spacer β-Cat T41A+1 | Rv | CTCTAAAACCGACAACTGCTCCCTCTTTGC |
| | | | 126 | scaffold | Fw | AGAGCTAGAAATAGCAAGTTAAAATAAGGCTAGTCCGTTATCAACTTGAAAAAGTGGGACCGAGTCG |
| | | | 127 | scaffold | Rv | GCACCGACTCGGTCCCACTTTTTCAAGTTGATAACGGACTAGCCTTATTTTAACTTGCTATTTCTAG |
| | | | 191 | R2 PBS-RT-V5 | Fw | GTGCTCCACTGCTCGGCCTGGATAGCACAGGTGGTGGCGGCGGAGGTATCCATTCCGGTGCAACTACAACTGCTCCCTC |
| | | | 192 | R2 PBS-RT-V5 | Rv | CGCGGAGGGAGCAGTTGTAGTTGCACCGGAATGGATACCTCCGCCGCCACCACCTGTGCTATCCAGGCCGAGCAGTGGA |
| | 1287 | pU6-TevQ1-β-Cat-WT-GFP11-Edit Fw | 220 | F Spacer β-Cat WT | Fw | CACCGTTGGCAGCAGCAGTCATATCGTTTT |
| | | | 221 | F Spacer β-Cat WT | Rv | CTCTAAAACGATATGACTGCTGCTGCCAAC |
| | | | 126 | scaffold | Fw | AGAGCTAGAAATAGCAAGTTAAAATAAGGCTAGTCCGTTATCAACTTGAAAAAGTGGGACCGAGTCG |
| | | | 127 | scaffold | Rv | GCACCGACTCGGTCCCACTTTTTCAAGTTGATAACGGACTAGCCTTATTTTAACTTGCTATTTCTAG |
| | | | 224 | F PBS-RT-GFP11-β-Cat WT | Fw | GTGCCAGCGGCGTTCACGTACTCGTGCAGCACCATGTGATCTCTTCCGCCGCCTCCGCCGAGATATGACTGCTGCTG |
| | | | 225 | F PBS-RT-GFP11-β-Cat WT | Rv | CGCGCAGCAGCAGTCATATCTCGGCGGAGGCGGCGGAAGAGATCACATGGTGCTGCACGAGTACGTGAACGCCGCTG |
| | 1288 | pU6-TevQ1-β-Cat-WT-GFP11-Edit Rv | 222 | R Spacer β-Cat WT | Fw | CACCGCAGTTGTCGTGGCACCGGAAGTTTT |
| | | | 223 | R Spacer β-Cat WT | Rv | CTCTAAAACTTCCGGTGCCACGACAACTGC |
| | | | 126 | scaffold | Fw | AGAGCTAGAAATAGCAAGTTAAAATAAGGCTAGTCCGTTATCAACTTGAAAAAGTGGGACCGAGTCG |
| | | | 127 | scaffold | Rv | GCACCGACTCGGTCCCACTTTTTCAAGTTGATAACGGACTAGCCTTATTTTAACTTGCTATTTCTAG |
| | | | 226 | R PBS-RT-GFP11 β-Cat WT | Fw | GTGCCGAGTACGTGAACGCCGCTGGAATTACAGGTGGTGGCGGCGGAGACTCTGGTATTCATTCCGGTGCCACGACA |
| | | | 227 | R PBS-RT-GFP11 β-Cat WT | Rv | CGCGTGTCGTGGCACCGGAATGAATACCAGAGTCTCCGCCGCCACCACCTGTAATTCCAGCGGCGTTCACGTACTCG |
| | 1293 | pU6-TevQ1-β-Cat-WT-SP-GFP11-Edit Fw | 124 | F Spacer β-Cat S33Y +3 | Fw | CACCGCAGCAGTCATATCTGGACTCGTTTT |
| | | | 125 | F Spacer β-Cat S33Y +3 | Rv | CTCTAAAACGAGTCCAGATATGACTGCTGC |
| | | | 126 | scaffold | Fw | AGAGCTAGAAATAGCAAGTTAAAATAAGGCTAGTCCGTTATCAACTTGAAAAAGTGGGACCGAGTCG |
| | | | 127 | scaffold | Rv | GCACCGACTCGGTCCCACTTTTTCAAGTTGATAACGGACTAGCCTTATTTTAACTTGCTATTTCTAG |
| | | | 200 | F PBS-RT WT-SP-GFP11 | Fw | GTGCATTCCAGCGGCGTTCACGTACTCGTGCAGCACCATGTGATCTCTTCCGCCGCCTCCGCCAGAATCCAGATATGACT |
| | | | 201 | F PBS-RT WT-SP-GFP11 | Rv | CGCGAGTCATATCTGGATTCTGGCGGAGGCGGCGGAAGAGATCACATGGTGCTGCACGAGTACGTGAACGCCGCTGGAAT |
| | 1294 | pU6-TevQ1-β-Cat-WT-SP-GFP11-Edit Rv | 160 | Spacer β-Cat T41A+1 | Fw | CACCGCAAAGAGGGAGCAGTTGTCGGTTTT |
| | | | 161 | Spacer β-Cat T41A+1 | Rv | CTCTAAAACCGACAACTGCTCCCTCTTTGC |
| | | | 126 | scaffold | Fw | AGAGCTAGAAATAGCAAGTTAAAATAAGGCTAGTCCGTTATCAACTTGAAAAAGTGGGACCGAGTCG |
| | | | 127 | scaffold | Rv | GCACCGACTCGGTCCCACTTTTTCAAGTTGATAACGGACTAGCCTTATTTTAACTTGCTATTTCTAG |
| | | | 185 | R2 PBS-RT-GFP11 S33Y | Fw | GTGCGTACGTGAACGCCGCTGGAATTACAGGTGGTGGCGGCGGAGGTATCCATTCCGGTGCAACTACAACTGCTCCCTC |
| | | | 186 | R2 PBS-RT-GFP11 S33Y | Rv | CGCGGAGGGAGCAGTTGTAGTTGCACCGGAATGGATACCTCCGCCGCCACCACCTGTAATTCCAGCGGCGTTCACGTAC |
| | 1238 | pTest-βCat-S33Y·mIRFP-670 | 1 | pCIG chbglob (Fw-Intron) | Fw | TTCGGCTTCTGGCGTGT |
| | | | 121 | β-Cat-BsmBI-int | Rv | TGAGACGCTCGTCTCTATAGTCCAGGTAAGACTGTT |
| | | | 122 | β-Cat-BsmBI-int | Fw | CTATAGAGACGAGCGTCTCAGGAATCCATTCTGGTGCCAC |
| | | | 123 | β-Cat-EcoRI | Rv | CCACTTGGCAGACCATCATC |
| | 1243 | pTest-βCat-S33Y-1 ×ALFA-tag-mIRFP-670 | 138 | 1× ALFA-tag | Fw | CTATGGCGGAGGCGGCGGACCCAGCAGACTGGAAGAAGAACTGAGAAGGCGCCTGACAGAGCCAGGTGGTGGCGGCGGA |
| | | | 139 | 1× ALFA-tag | Rv | TTCCTCCGCCGCCACCACCTGGCTCTGTCAGGCGCCTTCTCAGTTCTTCTTCCAGTCTGCTGGGTCCGCCGCCTCCGCC |
| | 1242 | pTest-βCat-S33Y-1 ×V5-tag-mIRFP-670 | 140 | 1× V5-tag | Fw | CTATGGCGGAGGCGGCGGAGGCAAGCCCATTCCAAATCCACTGCTCGGCCTGGATAGCACAGGTGGTGGCGGCGGA |
| | | | 141 | 1× V5-tag | Rv | TTCCTCCGCCGCCACCACCTGTGCTATCCAGGCCGAGCAGTGGATTTGGAATGGGCTTGCCTCCGCCGCCTCCGCCATAG |
| | 1241 | pTest-βCat-S33Y-1 ×GFP11-tag-mIRFP-670 | 170 | 1× GFP11-5G | Fw | CTATGGCGGAGGCGGCGGAAGAGATCACATGGTGCTGCACGAGTACGTGAACGCCGCTGGAATTACAGGTGGTGGCGGCGGA |
| | | | 171 | 1× GFP11-5G | Rv | TTCCTCCGCCGCCACCACCTGTAATTCCAGCGGCGTTCACGTACTCGTGCAGCACCATGTGATCTCTTCCGCCGCCTCCGCC |

| | | | | | | |
|---|---|---|---|---|---|---|
| **pTest-βCat Misexpression** | 1250 | pTest-βCat-S33Y-2 ×GFP11-tag-miRFP670 | DNA Fragment | 2×GFP11-tag | | AGCGTCTCACTATGGTGGCGGAGGCGGCAGAGATCACATGGTGCTGCACGAGTACGTGAACGCCGCTGGAATTACAGGAGGCGGC GGCGGACGTGATCATATGGTCCTTCACGAATATGTCAATGCCGCTGGGATCACCGGTGGTGGCGGCGGAGGAAAGAGACGAG |
| | 1246 | pTest-βCat-S33Y-3 ×ALFA-Tag-miRFP670 | DNA Fragment | 3×ALFA-Tag | | AGCGTCTCACTATGGTGGCGGAGGCGGCCCCAGCAGACTGGAAGAAGAACTGAGAAGGCGCCTGACAGAGCCAGGAGGCGGCGG CGGACCATCTAGGCTCGAGGAAGAACTGCGCAGAAGGCTGACTGAACCTGGCGGTGGCGGTGGCCCATCAAGGCTTGAAGAGGAA CTCAGACGCAGACTGACCGAGCCTGGTGGTGGCGGCGGAGGAAAGAGACGAG |
| | 1245 | pTest-βCat-S33Y-3 ×V5-tag-miRFP670 | DNA Fragment | 3×V5-tag | | AGCGTCTCACTATGGTGGCGGAGGCGGCAAGCCCATTCCAAATCCACTGCTCGGCCTGGATAGCACAGGAGGCGGCGGCGGAAAGC CAATACCTAATCCTCTGCTGGGACTCGACAGCACCGGCGGTGGCGGTGGCGATAAGCCTATTCCGAATCCGCTCCTGGGCCTCGATTCT ACTGGTGGTGGCGGCGGAGGAAAGAGACGAG |
| | 1244 | pTest-βCat-S33Y-3 ×GFP11-tag-miRFP670 | DNA Fragment | 3×GFP11-tag | | AGCGTCTCACTATGGTGGCGGAGGCGGCAGAGATCACATGGTGCTGCACGAGTACGTGAACGCCGCTGGAATTACAGGAGGCGGC GGCGGAAGGGACCACATGGTTCTCCATGAATATGTGAACGCTGCCGGCATCACTGGCGGTGGCGGTGGCCGTGATCATATGGTCCT TCACGAATATGTCAATGCCGCTGGGATCACCGGTGGTGGCGGCGGAGGAAAGAGACGAG |

| | | | | | | |
|---|---|---|---|---|---|---|
| **Misexpression** | 1187 | pCIG-human-β-CateninaS33Y | 38 | β-Cat EcoRI | Fw | AGCTGGAATTCTTTCTAACC |
| | | | 79 | β-Cat SmaI | Rv | TAGCCCGGGTCACAGGTCAGTATCAAACCAGG |

| | | | | | | |
|---|---|---|---|---|---|---|
| **Genomic amplification and sequencing** | | | 289 | Exon4 ggβ-Cat seq Fw | Fw | ttcaacgatttcttacag |
| | | | 290 | Exon4 ggβ-Cat seq Rv | Rv | ttgtagagaaggccttac |

**Supplementary Table 1. Oligonucleotides and synthetic DNA fragments used in this study.** Oligonucleotide sequences used for molecular cloning, base editing, prime editing, twin-prime editing, construct generation, and genomic amplification and sequencing are listed in the 5′–3′ orientation. Synthetic DNA fragments used to generate tandem-tag constructs are also included. Construct and oligonucleotide numbers correspond to the internal identifiers used in this study. For twin-prime editing epegRNAs, **F** and **R** designate the forward and reverse epegRNAs, respectively, whereas **Fw** and **Rv** in the *Direction* column indicate the complementary oligonucleotides used for cloning.
